# Quantifying mediation between omics layers and complex traits

**DOI:** 10.1101/2021.09.29.462396

**Authors:** Marie C. Sadler, Chiara Auwerx, Eleonora Porcu, Zoltán Kutalik

**Author notes:** Authors jointly supervised this work (Eleonora Porcu, Zoltán Kutalik).

## Abstract

**Background:** High-dimensional omics datasets provide valuable resources to determine the causal role of molecular traits in mediating the path from genotype to phenotype. Making use of quantitative trait loci (QTL) and genome-wide association studies (GWASs) summary statistics, we developed a multivariable Mendelian randomization (MVMR) framework to quantify the connectivity between three omics layers (DNA methylome (DNAm), transcriptome and proteome) and their cascading causal impact on complex traits and diseases.

**Results:** Evaluating 50 complex traits, we found that on average 37.8% (95% CI: [36.0%-39.5%]) of DNAm-to-trait effects were mediated through transcripts in the *cis*-region, while only 15.8% (95% CI: [11.9%-19.6%]) are mediated through proteins in *cis*. DNAm sites typically regulate multiple transcripts, and while found to predominantly decrease gene expression, this was only the case for 53.4% across ≈ 47,000 significant DNAm-transcript pairs. The average mediation proportion for transcript-to-trait effects through proteins (encoded for by the assessed transcript or located in *trans*) was estimated to be 5.27% (95%CI: [4.11%-6.43%]). Notable differences in the transcript and protein QTL architectures were detected with only 22% of protein levels being causally driven by their corresponding transcript levels. Several regulatory mechanisms were hypothesized including an example where cg10385390 (chr1:8’022’505) increases the risk of irritable bowel disease by reducing *PARK7* transcript and protein expression.

**Conclusions:** The proposed integrative framework identified putative causal chains through omics layers providing a powerful tool to map GWAS signals. Quantification of causal effects between successive layers indicated that molecular mechanisms can be more complex than what the central dogma of biology would suggest.

## Introduction

In the past decade, genome-wide association studies (GWASs) have identified thousands of genetic variants associated to complex traits [1], however mapping these variants to molecular processes and pathways still remains challenging [2]. A first step towards interpreting GWAS signals is to map trait-associated single nucleotide polymorphisms (SNPs) to genes. Naive approaches based on physical distance attribute SNPs to their closest gene [3] and many of them additionally take into account the linkage disequilibrium (LD) structure and GWAS association strengths (*i*.*e*. p-values) to compute gene scores [4, 5]. In a second step, scores from several genes can be combined and mapped to biological pathways by incorporating knowledge from external databases such as KEGG [6], Gene Ontology [7], WikiPathways [8], Reactome [9], or MSigDB [10], and making use of pathway enrichment analysis tools [5, 11, 12].

GWAS signals of common diseases predominantly fall into the non-coding genome [13] and both their enrichment in regulatory elements (e.g. quantitative trait loci (QTL) [13, 14]), as well as advances in omics technology [15], has motivated the establishment of large-scale consortia providing publicly available QTL datasets for molecular phenotypes such as DNA methylation (DNAm) [16], as well as transcript [17, 18], protein [19, 20, 21] or metabolite [22, 23] levels. Consequently, a next step in interpreting GWAS findings has been to integrate this new type of data, allowing to find diverse mediators of SNP-trait associations in a high-throughput, data-driven fashion. Integrative statistical methods combining GWAS and omics QTL summary data include colocalization tests [24, 25], summary versions of transcriptome-wide association studies (TWAS) [26, 27] and Mendelian randomization (MR) studies [28, 29]. Their application to a wide variety of GWAS datasets has resulted in the identification of many putative molecular trait-disease associations confirming known and highlighting potential new molecular mechanisms [30]. Colocalization methods identify shared QTL and GWAS signals, and while this might indicate causality between the molecular and GWAS trait, shared signals can also arise due to reverse causality (*i*.*e*. causal effect of the GWAS trait on the molecular trait [31]) or horizontal pleiotropy (*i*.*e*. the identified shared genetic variant drives the molecular and trait perturbation independently). In comparison, MR studies, which are conceptually similar to TWAS, that use multiple genetic variants as instrumental variables (IVs) are less prone to reverse causality and artefacts arising from LD patterns [32] - although horizontal pleiotropy can never be ruled out entirely. An important advantage of MR methods is that they allow the detection and elimination of pleiotropic markers. In addition, MR analyses allow the quantification - direction and magnitude - of the causal effect of the omic on the outcome trait.

With the advent of QTL datasets with increased sample sizes [16, 18], opportunities to integrate GWAS data with multiple molecular traits are no longer hampered by low statistical power. Previous efforts integrating multiple QTL omics data either adopted colocalization strategies [33, 34] or combined pairwise MR associations (two-step MR) [35, 36] to predict molecular mechanisms of the following scheme: omics trait 1 → omics trait 2 → outcome trait. While these approaches provide evidence for regulatory pathways, ascertaining their robustness can be difficult, since often only a single causal variant underlying these multiple associations was assessed [33, 35, 36]. As a consequence, the control over horizontal pleiotropy remained limited, although it was usually mitigated by the HEIDI (heterogeneity in dependent instruments) test statistic [28]. Furthermore, combining pairwise associations can lack the ability of inferring directionality between the different traits involved, an issue that can be identified by comparing the magnitude of QTL and GWAS effects [37]. Overall, while current integration methods test genetic downstream effects through omics traits, they often only accommodate the testing of a single molecular mediator.

Multivariable MR (MVMR) approaches have been proposed to identify multiple mediators of exposure-outcome relationships [38, 39]. These approaches enable the dissection of the total causal effect of an exposure on an outcome into a direct and indirect effect measured via mediators. Similar to MR, the use of genetic instruments allows for robust causal inference and MVMR has proven as an unbiased approach for mediation analyses, even in the presence of confounders [38, 39]. Hence, in addition to identifying causal effects through multiple layers, MVMR allows the quantification of mediation effects. Although not yet widely implemented on high-dimensional omics data, they provide great opportunities in the study of molecular mediation [40].

Here, we proposed a three-sample MVMR (3S-MVMR) framework to quantify the role of molecular mediators (omics trait 2) on a molecular exposure (omics trait 1) - complex trait relationship (Figure 1). We integrated methylomic, transcriptomic and proteomic QTL (mQTL, eQTL and pQTL, respectively) with GWAS summary data of 50 clinically relevant traits to perform mediation analyses and to estimate global mediation proportions (MPs). Three different combinations of exposure-mediator molecular traits were analysed: DNAm regulating transcripts in *cis*, DNAm regulating proteins in *cis*, and transcripts regulating their encoded protein in addition to proteins in *trans*. We performed simulation studies to estimate the bias of the defined MP under various parameter settings. In addition to quantifying the regulatory connectivity between each of these molecular layers, we investigated underlying factors driving high MPs, and hypothesized several mechanistic pathways between DNAm, gene expression and complex traits.

**Figure 1:**
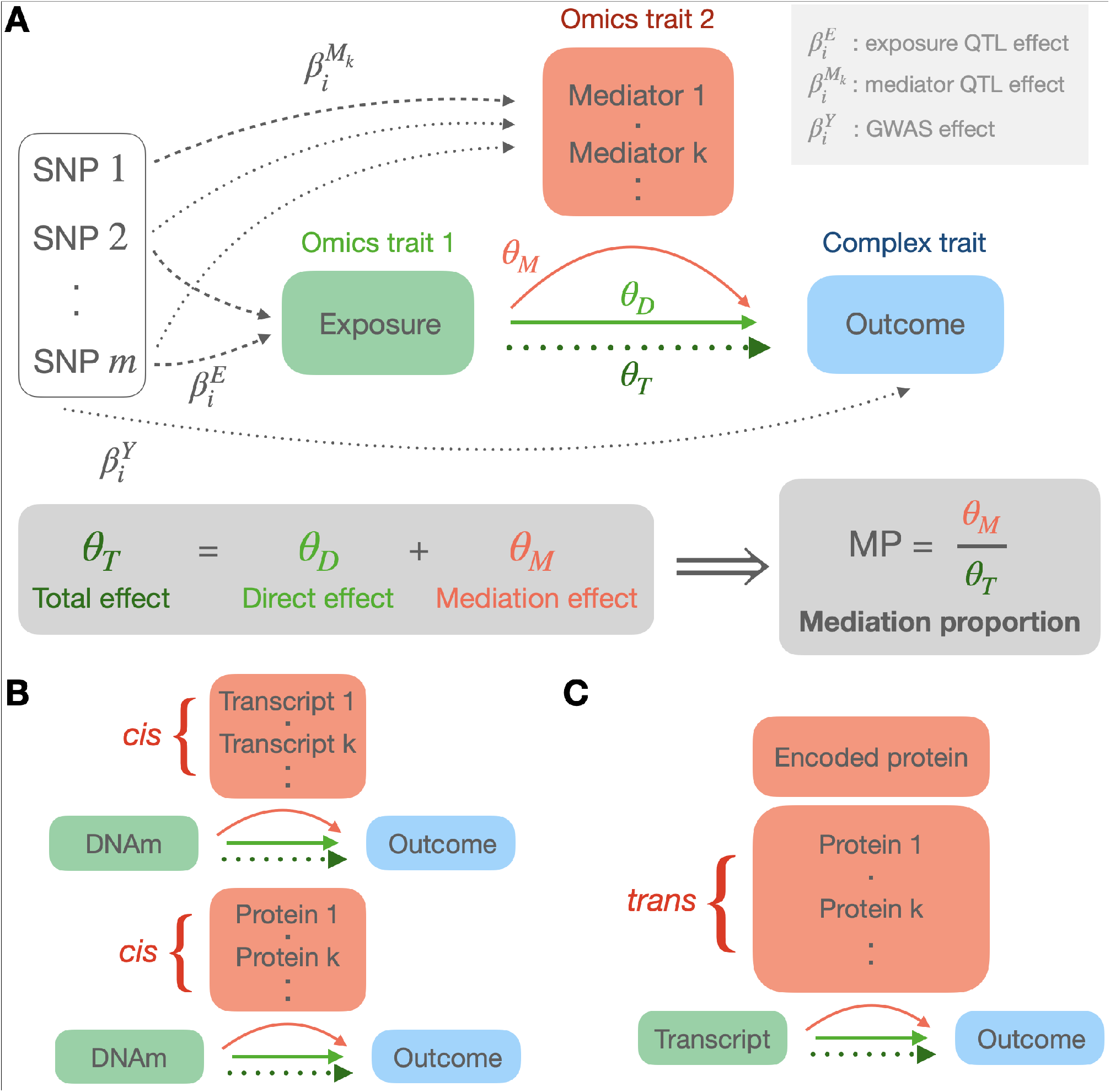
Overview of the MVMR design to quantify mediation of complex traits through DNAm, transcripts and proteins. **A)** General MVMR model: genetic instruments (SNPs) are selected to be directly associated (dashed arrow) with either the exposure (omics trait 1) or any mediator k (omics trait 2). The total effect *θ*_*T*_ (dotted arrow) of the exposure on the outcome (complex trait) is estimated in a univariable MR analysis based on exposure-associated SNPs only. The direct effect *θ*_*D*_ is estimated in a MVMR analysis on all valid instruments. The mediation effect *θ*_*M*_ results from the difference between *θ*_*T*_ and *θ*_*D*_, and allows to calculate the mediation proportion (MP). The genetic effect sizes *β* on the exposure, mediator and outcome come from m/e/pQTL and GWAS summary statistics, respectively. **B)** DNAm-to-complex trait effects were mediated once through transcripts in *cis* and once through proteins in *cis*. **C)** Transcript-to-complex trait effects were mediated through the protein the transcript is encoding for (encoded protein; if available in the dataset), as well as through proteins in *trans*. Except for the encoded protein, mediators were required to be causally associated to the exposure in both DNAm- and transcript-exposure settings.

## Results

### Overview of the method

We performed univariable and multivariable MR to estimate total and direct effects, 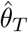 and 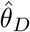, respectively, of molecular exposures on 50 outcomes through various molecular mediators (Figure 1; Equation 1 and 3). MP estimates were then calculated as the ratio of the indirect effect through the molecular mediators to the total effect of the exposure on the outcome trait [41] and computed only for exposure-outcome pairs with significant Bonferroni-corrected 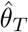 effects, grouped by trait, trait category and all pairs combined. We further filtered out exposure-outcome pairs whose exposures have no significant causal effect on any potential mediator as the link between those pairs cannot be mediated. For completeness, however, we also present results for the scenario when the last filtering step is omitted.

While weak genetic instruments in univariable MR analyses can introduce a bias towards the null [42], it has been shown that this bias can be in any direction in MVMR studies [43]. Both the sample size and the choice of instruments and mediators can contribute to biases in various directions [43], leading to under- or over-estimations of the MP. To quantify this bias and assess the sensitivity and robustness of estimated 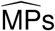, we conducted simulation studies mimicking the settings that emerge in real data applications for either DNAm or transcript levels as exposure (Methods) (Figure S1).

### Simulation results

Simulations showed that the bias in the estimated MPs 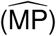 is minimal for the settings most relevant for real data we explored (Figures S2-3; Table S2; Methods). A determining factor in accurately estimating MPs was the sample size of the mediator QTL effects. Low sample sizes resulted in significant underestimations of the MP, with sample sizes of 3,000 compared to 30,000 resulting in a 20% relative decrease (6% in absolute values) of the estimated 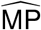 in the DNAm-exposure simulation settings (Figure 2A). The reason for this significant underestimation was the omission of relevant mediators with 0.45/3 (15%) being missed at a sample size of 3,000 in the DNAm-exposure simulation settings (Figure 2B). We further tested the robustness of the 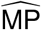 with respect to the number of included mediators by varying the mediator selection threshold P_EM_ (Methods). At more stringent threshold, relevant mediators were more likely to be missed resulting in an underestimation of the 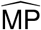 (Figure S4). Importantly, including irrelevant mediators at more lenient thresholds did not bias the 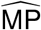, although a critical point was reached upon the inclusion of > 10 irrelevant mediators where the estimated 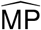 started to become underestimated in the transcript-exposure setting (Figure S4). The used transcript and DNAm QTL datasets provide SNP effect sizes in *cis* of the assessed transcript and probe, respectively, and were primarily restricted to significant mQTLs for the latter. Thus, SNP-exposure effects for SNPs serving as mediator instruments are often missed and set to zero. However, our simulation studies, which mimicked this scenario by setting non-significant effects to zero (Methods), showed that this did not induce any bias.

**Figure 2:**
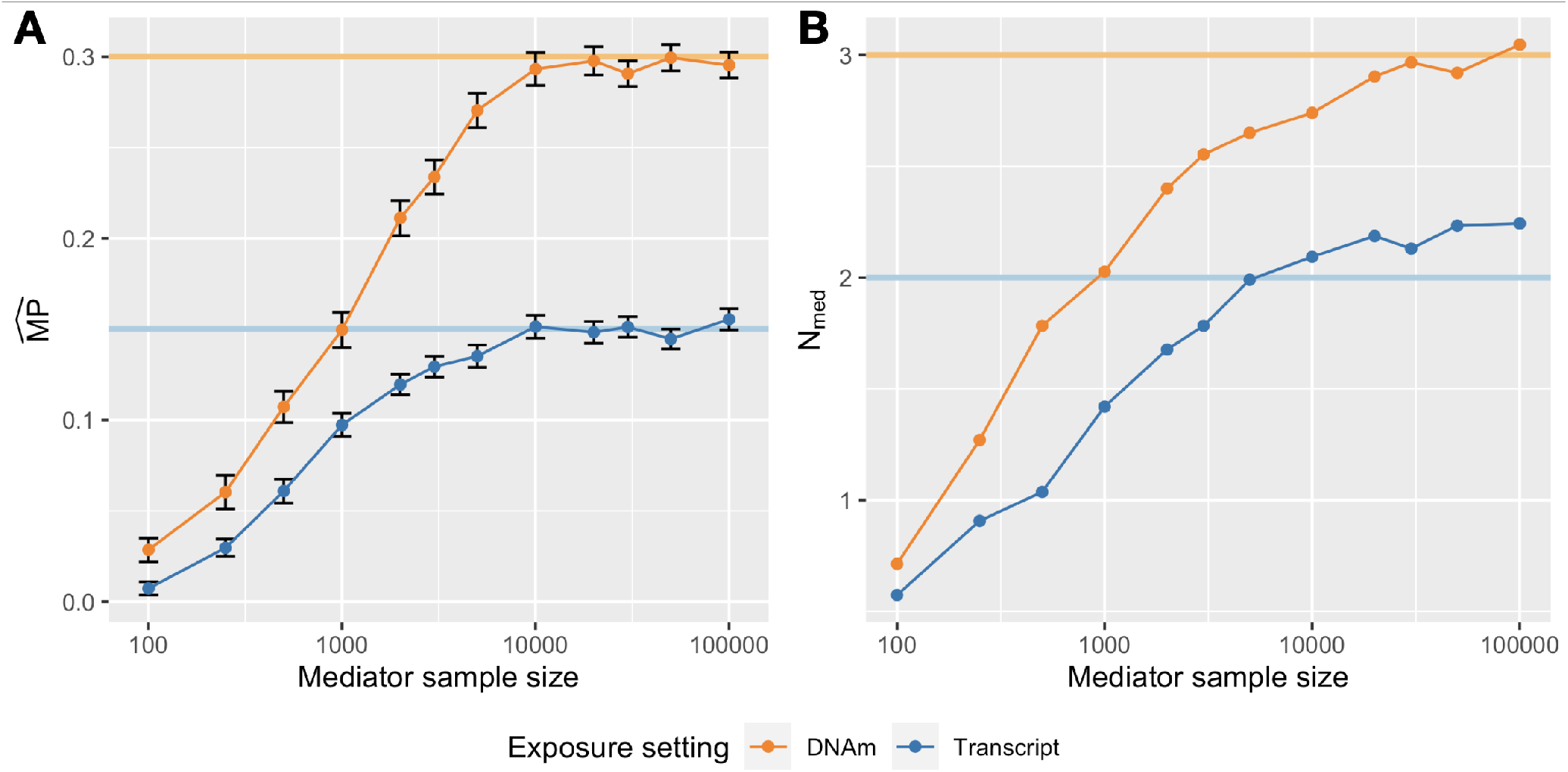
Simulation results in DNAm- (orange) and transcript- (blue) exposure settings to assess the impact of the mediator sample size on the A) estimated 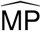 and B) number of selected mediators. For a given mediator sample size, 300 exposure-outcome pairs were simulated on which an 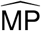 and 95% CI (error bars) were estimated. The true MP of the model was 0.3 and 0.15, and the true number of relevant mediators was 3 and 2 in the DNAm-exposure and transcript-exposure setting, respectively, as indicated by the solid horizontal lines.

### DNAm-to-complex trait effects mediated by gene expression in *cis*

Across 50 traits (Table S1), we evaluated the mediation of 2,069 DNAm-trait causal pairs by transcripts in *cis*. The 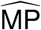 for each of the 41 traits influenced by at least 10 DNAm probes ranged from 18.0 to 78.0% (mean: 36.9%, 95% CI: [13.5%-60.3%]) (Figure 3A). Regressing 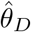 against 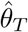 for all pairs combined and accounting for regression dilution bias (Equation 4) yielded an 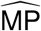 of 37.8% (95% CI: [36.0%-39.5%]) (Figure 3B). Grouping the traits into 10 physiological categories (Table S1) showed that the 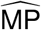 was highest for hepatic biomarkers (mean: 46.6%, 95%CI: [41.5%-51.7%]), followed by renal biomarkers (mean: 43.5%, 95%CI: [37.5%-49.5%]). In contrast, adiposity-related and hormonal traits exhibited the lowest 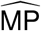 (Figure 3B, Figure S5).

**Figure 3:**
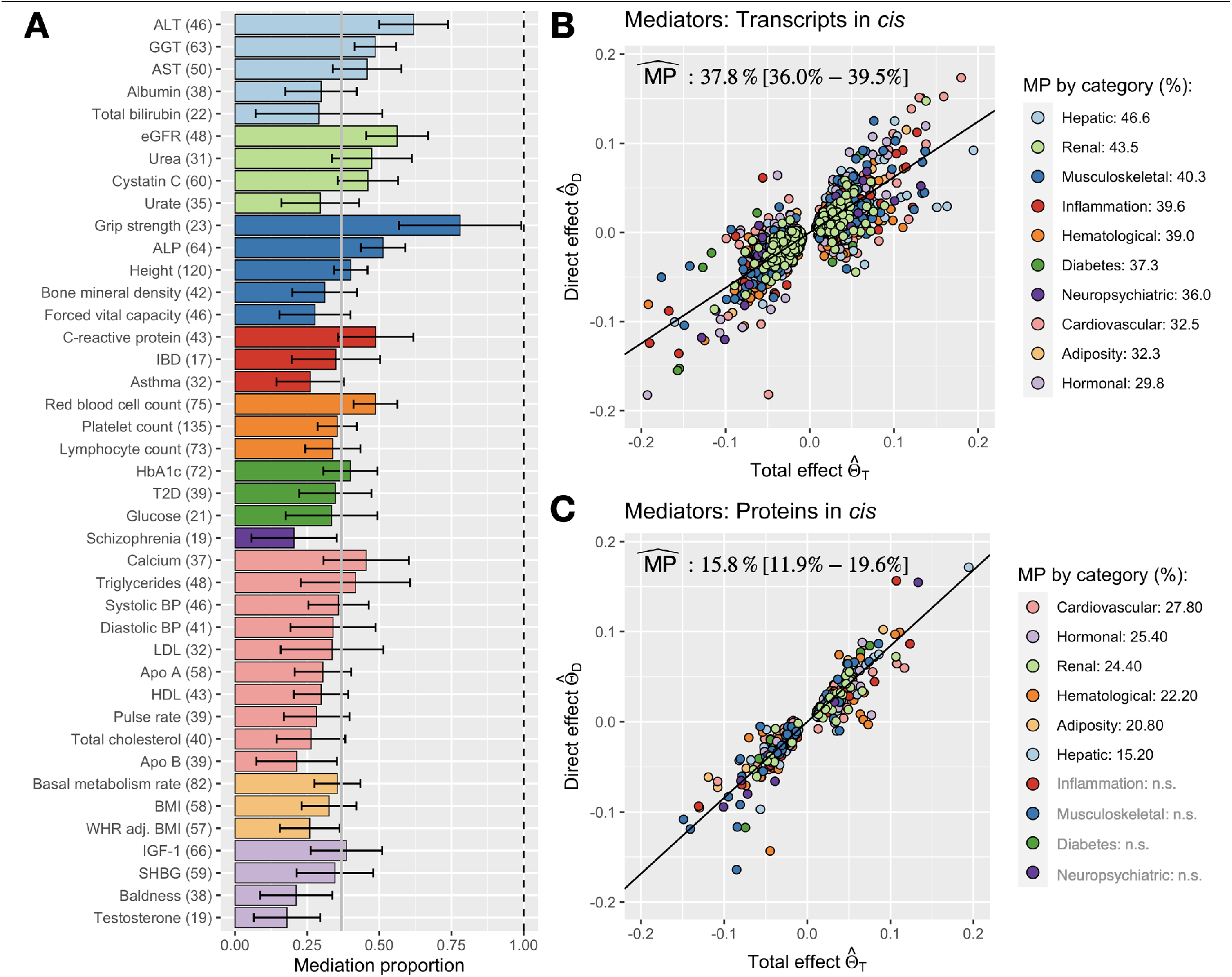
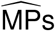 for transcripts and proteins in *cis* mediating DNAm-to-trait effects. **A)** 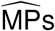 by trait in the DNAm-to-trait via transcripts in *cis* analysis. Error bars denote the 95% CI, and the grey vertical bar shows the mean 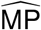 across the traits. Only traits with ≥ 10 DNAm-trait pairs are displayed (41 traits with the exact number of evaluated pairs indicated in parentheses), colour-coded by their physiological category as defined in the legends of B) and C). **B)** All DNAm-trait pairs with traits being grouped into 10 physiological categories. The global 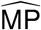 in % with 95% CI is shown in the plotting area and individual category 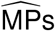 in the legend. **C)** Same analysis as in B), but with mediators being proteins in *cis*. Category 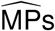 not significantly different from zero (n.s.) are written in grey.

The average number of mediator transcripts was 3.3 per methylation-trait pair, indicating that the impact of methylation is not mediated by a single transcript. To further explore this observation, we assessed the extent to which DNAm→trait effects were mediated by the single most significantly DNAm-associated transcript (“top” transcript; Methods), as opposed to all transcripts in *cis*. This resulted in an 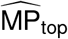 of 26.0% (range: [13.0%-46.8%]) averaged across the 41 traits, and an 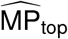 of 26.6% (95% CI: [25.1%-28.1%]) when aggregating the 2,069 DNAm-trait pairs. This significant drop in the 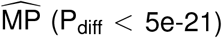 corroborates our initial hypothesis that DNAm sites regulate the expression of multiple transcripts in the *cis* region.

Including DNAm-trait pairs with testable transcripts in the *cis* region, but not causally linked to the assessed DNAm site (2,623 DNAm-trait pairs, Methods: adjusted MP calculation; Table S3), decreased the overall 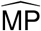 to 28.3% (95% CI: [26.9%-29.8%]) (Figure S7). While it may seem to be a more objective measure of the importance of the transcriptome in mediating DNAm-to-phenotype effects, it is overly conservative since the set of testable transcript mediators (N = 19,250 [18]) is a magnitude lower than that of the whole transcriptome [44]. A distribution of the number of times no mediation analysis could be conducted due to the absence of (causally associated) transcripts in the region or insufficient (exposure-associated) IVs is shown in Figure S24.

### Transcripts levels are under tighter DNAm control than protein levels

Next, we investigated the role of protein levels as mediators. Assessing the same DNAm-trait pairs as previously, we performed mediation analyses based on a potential mediator set of 2,838 proteins in total (INTERVAL pQTL dataset [19]). The estimated 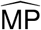 equalled 15.8% (95% CI: [11.9%-19.6%]) across 328 DNAm-trait pairs with at least 1 mediator protein (Figure 2C). Highest 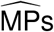 were obtained for cardiovascular traits (mean: 27.8%, 95% CI: [19.9%-35.8%]) (Figure S6). Given the lower sample size of the pQTL dataset and the results from the simulation studies, a drop in MP was expected. Not only the lower sample size, but also the lower number of testable proteins contributed to this decrease. To compare the difference in MP due to the mediators being transcripts instead of proteins, we repeated the analysis on the common set of transcripts and their encoded proteins (N = 2,145). We observed a drop in the adjusted 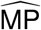 from 28.3% (95% CI: [26.9%-29.8%], 2,623 DNAm-trait pairs) to 8.15% (95% CI: [7.11%-9.19%], 2,111 DNAm-trait pairs) for transcripts and from 1.24% (95%CI: [0.66%-1.83%], 2,380 DNAm-trait pairs) to 0.85% (95%CI: [0.30%-1.39%], 2,111 DNAm-trait pairs) for proteins (Figure S8). A key difference in the two mediation analyses was the number of mediators (*N*_*med*_) found to be causally associated to the DNAm site and subsequently included in the mediation analysis (mean *N*_*med,transcript*_ = 0.48 and mean *N*_*med,protein*_ = 0.12). Restricting MP calculations to the same DNAm-trait pairs with at least one transcript and protein mediator, no statistical difference between the two MPs could be detected (P_diff_ = 0.28; Figure S9). Besides differences in sample size, a previous pQTL study of larger sample size (N = 30,931) reported strong differences in the underlying genetic architecture of transcript and protein levels, with less than a third of pQTLs being also eQTLs [21]. Accordingly, we found that only 333 out of 1,510 transcripts (*i*.*e*. those with a corresponding protein product present in the INTERVAL dataset and having at least 3 independent eQTLs to be used as IVs) could explain the levels of their encoded protein at a nominal significance threshold (Methods; Table S6). Focusing on DNAm-trait pairs where the top transcript mediator was the same than the top protein mediator (N = 106), the proportion of protein levels causally linked to their transcript levels increased to 72%. While both mediation analyses yielded similar results for transcript and protein levels with the same QTL structure, the findings suggest that overall, the genetic architecture of mQTLs is more similar to the one of eQTLs than to the one of pQTLs, which translates to a stronger DNAm-trait mediation through transcripts than through protein levels.

### Determining factors of mediation proportions

We further explored underlying factors driving high MPs through transcript levels (Figure 4A). 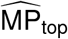 decreased with increased distances between the DNAm site and the gene transcription start site (TSS) of the top transcript (*ρ* = −0.076, P = 5.2e-4; Figure 4B). Further investigations revealed that this distance is negatively correlated to the DNAm-to-transcript MR squared effect size, 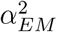, (*ρ* = −0.13, P = 3.1e-19; Figure 4C), which in turn is a good predictor for high MPs (*ρ* = 0.39, P = 2.5e-75; Figure 4D). The mediation proportion was the highest for DNAm sites residing in the first exon, followed by those in the 5’UTR, within 200bp of the TSS and finally lowest for those within 1500bp and in the gene body (Figure S10).

**Figure 4:**
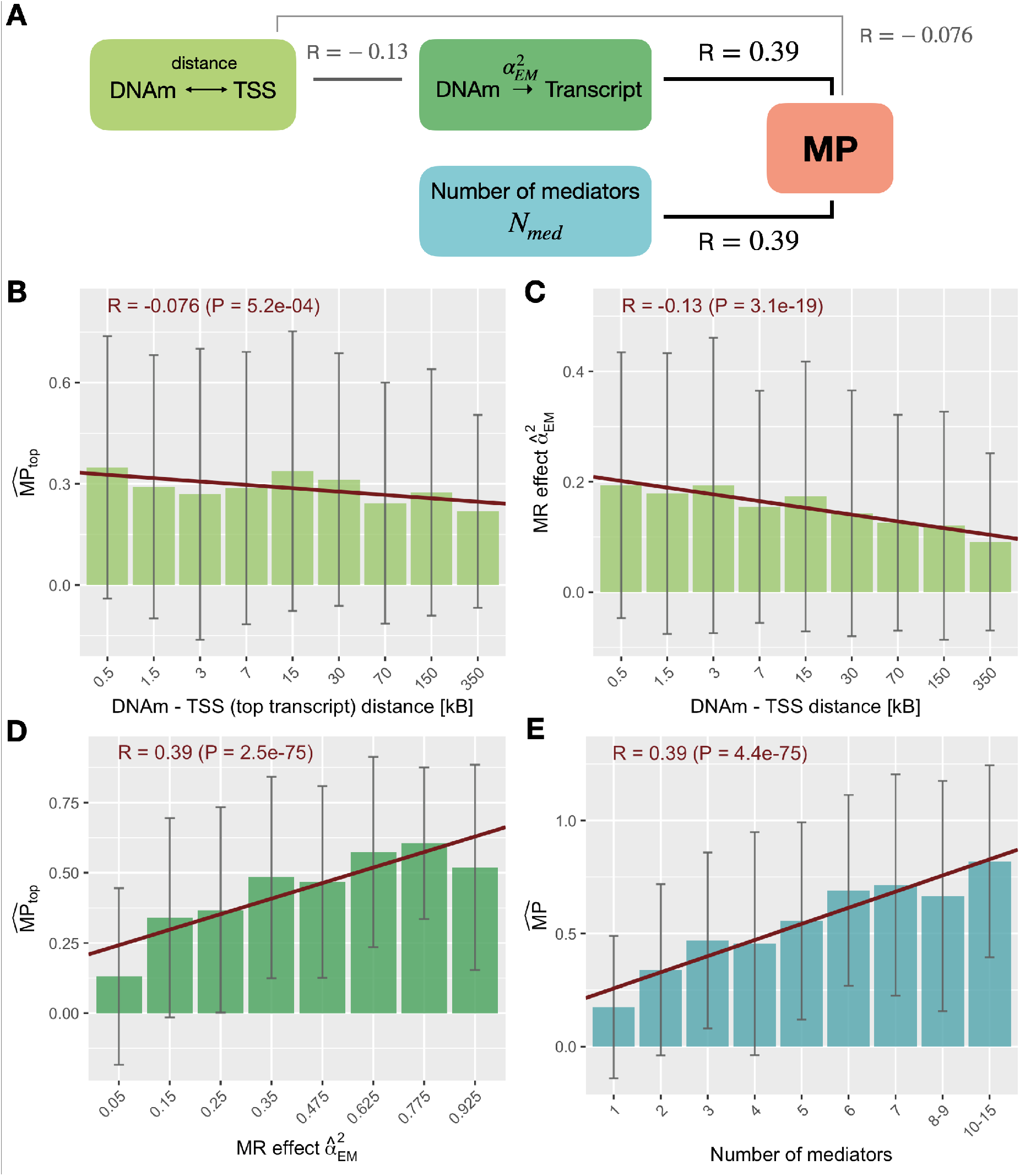
Exposure-to-mediator regulatory strength and number of mediators explaining MPs. **A)** Summary of the correlations (R) between MP and DNAm-to-transcript causal MR effects 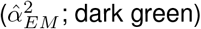, distance between the DNAm site and transcription start site (TSS; light green) and number of mediators (*N*_*med*_; blue). **B)** Average 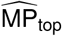 of DNAm-transcript pairs stratified according to the distance between the DNAm site and the TSS of the top transcript. All DNAm-trait pairs with at least one mediator were included. **C)** Average MR causal effects 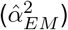 of DNAm-transcript pairs stratified according to the distance between the DNAm site and the TSS. Unique DNAm-transcript mediator pairs across all DNAm-trait pairs were included. **D)** Average top 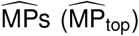 of DNAm-trait pairs stratified according to DNAm-to-top transcript MR causal effect size 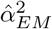. All DNAm-trait pairs with at least one mediator were included. **E)** Average 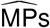 of DNAm-trait pairs stratified according to the number of mediators. All DNAm-trait pairs with at least one mediator were included. In every calculation, Pearson correlations and corresponding p-values (P) between the two respective quantities were calculated on DNAm-trait/DNAm-transcript pairs prior stratification. Error bars represent standard deviations and the red slope represents the regression fit between the bin’s positions and heights, and serves merely for visualization purposes.

DNAm inhibiting the binding of transcription factors (TFs) and thus repressing gene expression is often alluded to as the classical mechanism of action for DNAm [45] and might be driving this observation. However, many other mechanisms have been hypothesized [46] and many might still be unknown [47]. From the 1,066,307 unique DNAm-to-transcript causal effects assessed, 47,445 were significant at P < 4.7e-8. Although negative effects had a larger magnitude than positive ones (two-sided t-test: P = 0.0082) only 53.4% of DNAm→transcript causal effects were negative. Stratifying DNAm sites with respect to their location on the assessed transcript, we found that DNAm sites situated in the first exon and nearby the TSS were enriched for negative effects (P = 2.7e-3, 1.2e-5 and 3.8e-4 for 1st exon, TSS1500 and TSS200, respectively), whereas those in the gene body were enriched for positive ones (P = 2.2e-10; Table S4). These observations are in line with previous studies that only showed a slight trend for negative methylation-gene expression correlations [46, 48, 49, 47]. We further tested whether the MR DNAm-to-transcript causal effects correlated with reported methylation-transcript correlations [48] and found a strong agreement (*ρ* = 0.39, P = 2.6e-18, 471 DNAm-transcript pairs).

Consistent with higher MPs when mediating through multiple transcripts, we found a strong correlation between the number of mediators and the MP (*ρ* = 0.39, P = 4.4e-75; Figure 4E). Many of these mediators were correlated amongst each other, which in theory should be accounted for via the multivariable Mendelian randomisation. To ensure that this was the case, we repeated the mediation analysis with uncorrelated mediators (R_med_ < 0.3; Methods). The mean number of selected mediators dropped by more than half, from 3.3 to 1.2 (Figure S11), and the 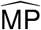 across all the DNAm-trait pairs decreased 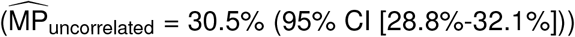, while remaining significantly higher than 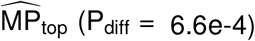. Decreasing the R_med_ threshold to 0.2 and 0.1 did not significantly decrease 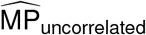 (P_diff_ > 0.05), which stabilized at 29.2% (95% CI: [27.5%-30.8%] for R_med_ < 0.1 (Figure S11).

Finally, we assessed the influence of the p-value threshold P_EM_ to select mediators based on the exposure-to-mediator causal effect (default P_EM_ = 0.01 for which N = 2,069 DNAm-trait pairs with at least 1 mediator were found). With a more lenient threshold (P_EM_ = 0.05), more DNAm-trait pairs with mediators emerged (N = 2,189). Conversely, with a more stringent threshold (P_EM_ = 0.001), less pairs were detected (N = 1,881). No differences in MPs between the three settings were found (P_diff_ > 0.05; Figure S12), but when calculating the adjusted MP (inclusion of all DNAm-trait pairs with potential transcript mediators in the *cis*-region) on a common set of DNAm-trait pairs (N = 2,543, 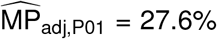 (95% CI: [26.1%-29.2%])), a significantly higher MP for the more lenient threshold 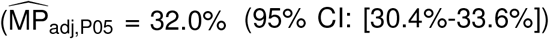; P_diff_ = 1.1e-4), and significantly lower MP for the more stringent threshold were observed 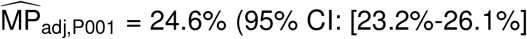; P_diff_ = 4.8e-3; Figure S13). Overall, these sensitivity analyses showed that the estimated MPs remain robust with respect to correlations within the selected mediator set, while also suggesting that the P_EM_ may be threshold-sensitive and mediators selected at the 0.01 p-value threshold may lead to conservative MP estimates.

### Transcript-to-complex trait effects mediated by proteins

Next, we quantified the role of proteins in mediating transcript-to-trait causal effects. First, we identified 3,848 significant transcript-trait pairs (P_T_ < 5e-5, ≥ 5 IVs; Table S3) across the 50 traits and performed mediation analyses through the protein encoded by the transcript (if present in the INTERVAL pQTL dataset [19]), in addition to any other protein in *trans* (*i*.*e*. any protein which is not encoded by the investigated transcript) causally associated to the transcript (P_EM_ = 1e-3; Figure 5A). The estimated 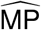 for the 1,577 transcript-trait pairs with at least 1 mediator was 5.27% (95%CI: [4.11%-6.43%]) and significantly higher than average for cardiovascular traits (P_diff_ = 8.7e-3; mean = 9.62%, 95% CI: [6.58%-12.7%]; Figure 5B). A distribution of the number of times transcript-trait pairs could be assessed in a mediation analysis through proteins is shown in Figure S25. When further restricting the mediation analysis to only those transcript-trait pairs for which the encoded protein was present, we observed an 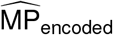 of 5.08% (95% CI: [2.62%-7.53%], 333 transcript-trait pairs) which increased, albeit not significantly, when additionally considering mediator proteins in *trans* (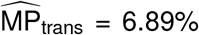; 95% CI: [4.17%-9.61%]; Figure S14). As mentioned previously, less than a quarter of the causal MR effects of transcripts on their encoded proteins were nominally significant and since we found exposure-mediator effects to be driving the MP (Figure 4B), we next focused on the 93 transcript-trait pairs for which the encoded protein was nominally significantly associated to the transcript (Figure S16). We observed a non-significant increase in the 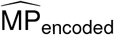 to 13.7% (95% CI: [4.34%-23.0%]) and in the 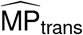 to 16.6% (95% CI: [8.19%-25.1%]). Stratifying traits by broad categories (e.g. metabolite, protein, physical measurement; Table S1), highest MPs were achieved for protein outcome traits (e.g. apolipoprotein B, alkaline phosphatase; Figure S15 and S17).

**Figure 5:**
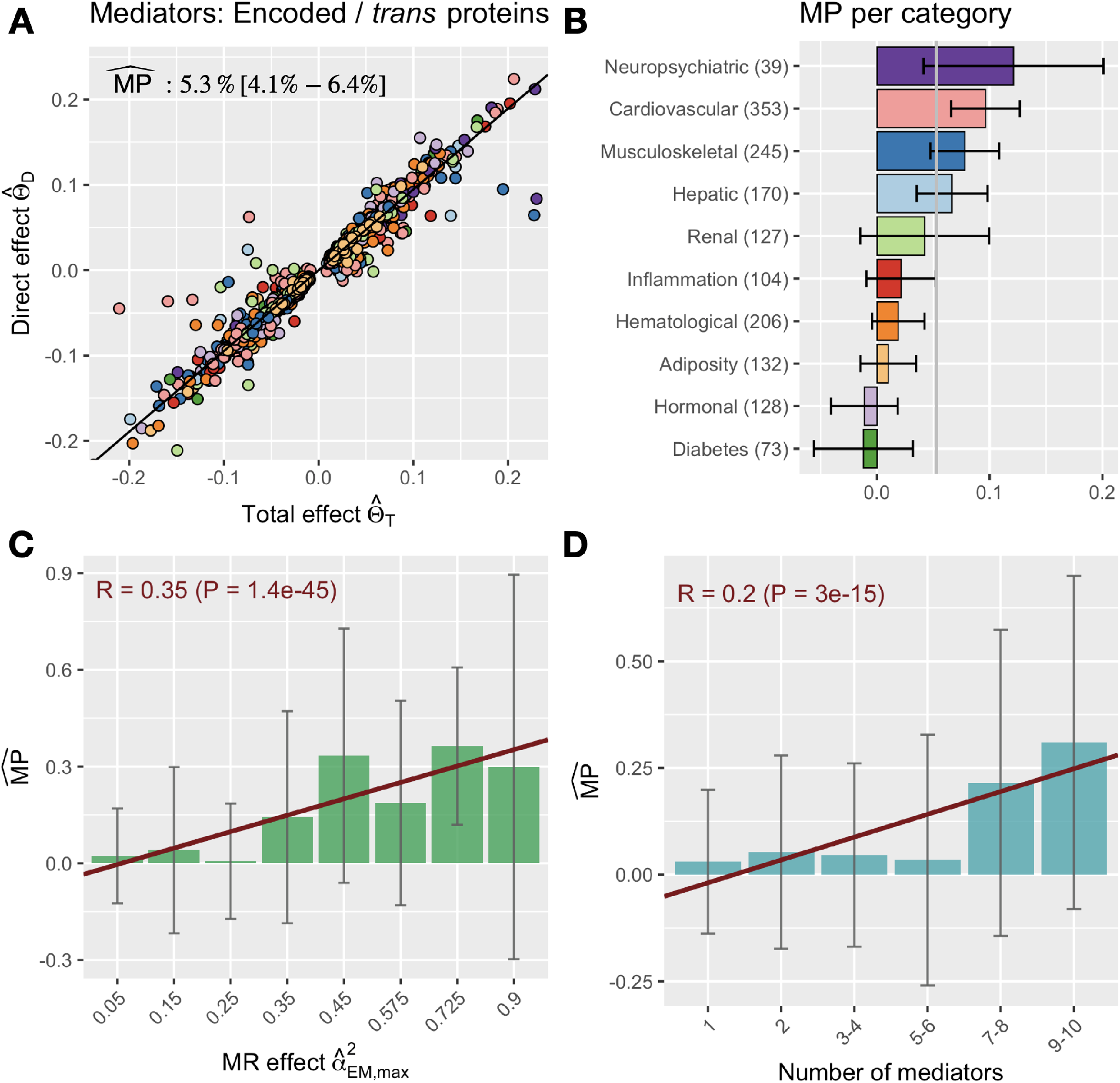
Proteins mediating transcript-to-trait causal effects. **A)** All transcript-trait pairs with with at least one mediating protein (encoded or/and in *trans*). Pairs are colour-coded by physiological categories as defined in B) and overall 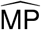 in % with 95% CI is shown. **B)** 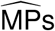 by trait category for the same pairs. Error bars denote the 95% CI, and the grey vertical line shows the mean 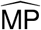 across transcript-trait pairs. The number of evaluated transcript-trait pairs in each category is indicated in parentheses. **C)** 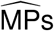 of transcript-trait pairs stratified according to the maximum transcript-to-protein MR causal effect size within the set of included mediator proteins. **D)** 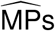 of transcript-trait pairs stratified according to the number of mediators. Correlations (R) and corresponding p-values (P) in C) and D) were calculated on transcript-trait pairs (same ones as in A) and B)) prior to stratification. Error bars represent standard deviations and the red slope represents the regression fit between the bin’s positions and heights, and serves merely for visualization purposes.

As for the DNAm-trait mediation analysis, we confirmed that strong exposure-mediator effect size 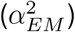 was the major driver of high MPs (Figure 5C). We calculated the correlation between MP and the maximum causal effect size squared between the transcript and any of its mediator proteins 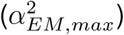, yielding *ρ* = 0.35 (P = 1.4e-45, N_pairs_ = 1,577, mean *N*_*med*_ = 2.15). Additionally, we found a significant correlation between *N*_*med*_ and MP (*ρ* = 0.20, P = 3.0e-15, N_pairs_ = 1,577). We performed further sensitivity analyses to assess the influence of the P_EM_ threshold (default P_EM_ = 1e-3). Considering all transcript-trait pairs (including those with no encoded protein), choosing a more lenient threshold (P_EM_ = 0.01) resulted in more transcript-trait pairs to be evaluated (N_pairs_ = 2,820), but not in a significant change in MP (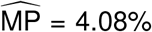, 95% CI: [3.14%-5.03%]; P_diff_ > 0.05; Figure S18). On the other hand, a more stringent threshold (P_EM_ = 1e-4) resulted in fewer transcript-trait pairs (N_pairs_ = 758) and a significantly higher MP (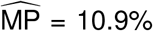, 95% CI: [9.02%-12.9%]; P_diff_ = 7.2e-7), as consequence of selecting transcript-trait pairs with higher 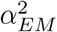 (Figure S21).

Finally, we aimed at validating our results using a different protein dataset. While there are publicly available pQTL datasets of larger sample size, the number of tested proteins in these studies is orders of magnitude lower (e.g. 71 proteins (N = 6,861) in the Framingham Heart Study [20]; 90 proteins (N = 30,000) in the SCALLOP consortium [21]). We used pQTL summary statistics of 41 cardiovascular proteins released by the SCALLOP Consortium [21] which overlapped with our main pQTL dataset from the INTERVAL Consortium [19], as well as with our main eQTL dataset of the corresponding transcripts (eQTLGen Consortium [18]). In a first step, we tested the agreement of MR causal effects of the transcripts on their encoded protein between the two protein datasets. Among the 38 tested transcript-protein pairs (≥ 1 eQTL), we observed a very strong correlation of causal effect sizes (*ρ* = 0.74, P = 9.4e-8) and no difference in their magnitude (two-sided t-test: P > 0.05). Given the larger sample size of the SCALLOP dataset, standard errors of the effect estimates were on average three times smaller (Figure S19). Due to the small number of overlap between the proteins and transcripts in the three datasets, there were only 6 transcript-trait pairs that could be compared in the mediation analysis through encoded protein, and for these, there was no significant difference in the direct effects 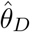 (Figure S20). Overall, while we could replicate causal effect sizes between the transcript and protein levels, the small number of available proteins did not allow to reliably quantify the bias in our estimated MP caused by the comparably small sample size of the pQTL dataset.

### DNAm-to-complex traits mechanisms of action

In addition to providing insights into global patterns governing the mediation between different intermediate phenotypic layers and functional traits, our analyses generated plausible hypotheses regarding specific biological pathways. Recently, the involvement of the anti-oxidant and anti-inflammatory protein PARK7 in inflammatory bowel disease (IBD) has been brought to light [50, 51, 52, 53]. While the exact role of the protein in the disease remains debated, reduced intestinal expression of *PARK7* was observed in patients and mouse models for IBD [53]. Moreover, *Park7* knockout mice were shown to have increased levels of pro-colitis bacterial species in their microbiome [54, 52] and experience aggravated symptoms of experimental-induced colitis [53]. In line with these observations, DNAm of the *PARK7* promoter probe cg10385390 (chr1:8’022’505) decreased both *PARK7* transcript (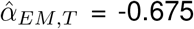, P = 2.7e-4) and protein (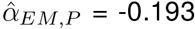, P = 2.0e-3) expression (Figure 6A). High transcript (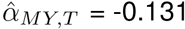, P = 1.7e-7) and protein (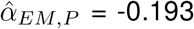, P = 0.31) levels decrease IBD risk, resulting in an overall increased IBD risk upon DNAm 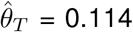. Interestingly, early GWAS identified the region as a susceptibility locus for IBD, listing *TNFRSF9* as the top candidate gene [55, 56], thereby exemplifying how the integration of multiple omics layers can help to identify further causal genes.

**Figure 6:**
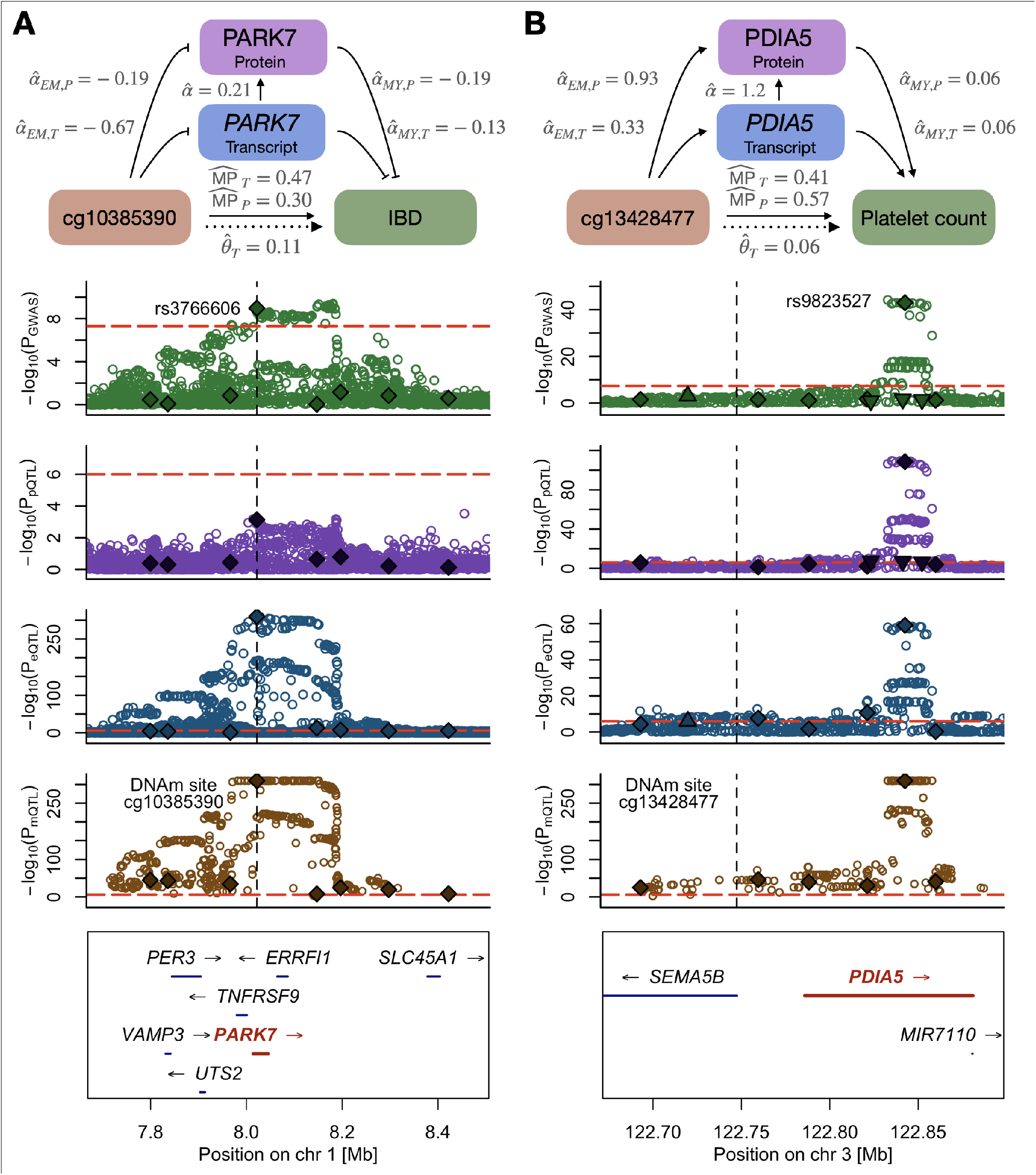
Plausible DNAm-transcript/protein-trait regulatory mechanisms between **A)** *PARK7* and irritable bowel disease (IBD) and **B)** *PDIA5* and platelet count. The top row displays a schematic of the mechanism with calculated univariable and multivariable MR effects. The four following rows show the regional SNP associations (-log_10_(p-values)) with the trait (green), encoded protein (purple), transcript (blue) and DNAm (brown) probe, respectively. Solid diamonds represent DNAm-associated instruments used in the univariable (for 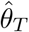 calculation) and multivariable (for 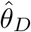 calculation) MR analyses. Upwards and downwards pointing triangles are transcript- and protein-associated SNPs, respectively, that were additionally included in the MVMR instrument set. Red dashed lines indicate the significance thresholds of the respective SNP associations and the vertical black dashed line represents the DNAm probe position. Bottom row illustrates the positions and strand direction of the genes in the locus.

Despite often being associated with decreased expression [45], our data provides examples of methylation boosting expression. For instance, DNAm of cg13428477 (chr3:122’748’086) increased *PDIA5* expression 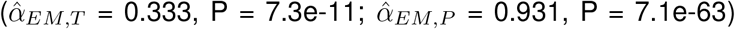, whose levels subsequently increased platelet count 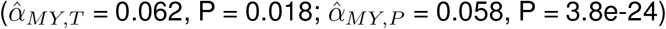, so that DNAm resulted in increased platelet count 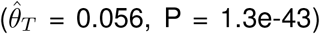 (Figure 6B). Association between the *PDIA5* locus and platelet count was reported through GWAS [57]. Platelets are small cell fragments produced by megakaryocytes, which themselves are derived from hematopoietic stem cells. Accordingly, *PDIA5* has a binding site for the hematopoietic stem and progenitor cell TF MEIS1 [58] and is overexpressed in megakaryocytes as compared to other blood cell types [59]. Further studies showed that *pdia5* protein knockdown in zebrafish resulted in strongly decreased platelet count [60], matching our findings and confirming the role of *PDIA5* in thrombopoiesis. Additional putative regulatory mechanisms of DNAm-to-complex traits through transcript and protein levels are shown in Table S8-9, respectively, and were selected based on 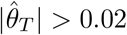 and 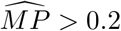.

While the aforementioned DNAm → gene expression → trait mechanisms were supported by both differential transcript and protein levels, other examples for which protein expression could not be assessed due to lack of pQTL data still reflect highly plausible mechanisms. For instance, we observed that DNAm of cg09070378 (chr1:161’183’762) decreased asthma risk (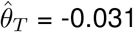, P = 8.1e-11) by reducing *FCER1G* expression (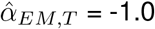, P = 3.5e-18), a gene listed in the KEGG pathway for asthma (hsa05310) and whose expression associated with an increased risk for asthma (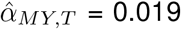, P = 3e-12) (Figure S21). The *FCER1G* promoter was found to be hypomethylated in patients with atopic dermatitis, with DNAm levels correlating negatively with the gene’s expression [61], suggesting a broad role of *FCER1G* in allergic disorders. Our data also supports and provides a mechanistical explanation for the recent finding that reduced *IFNAR2* expression causally decreases the odds of severe coronavirus disease 2019 (COVID-19) [62, 63], which was later supported by the increased susceptibility for severe COVID-19 in individuals with rare loss-of-function mutations in *IFNAR2* [64]. Indeed, we found that DNAm of the *IFNAR2* promoter probe cg13208562 (chr21:34’603’264) decreased the gene’s expression (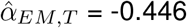, P = 2.4e-19) (Figure S22). As *IFNAR2* expression protects against hospitalization following COVID-19 infection (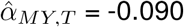, P = 4.2e-6), DNAm of the locus increased the risk of severe infection (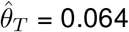, P = 8.5e-13).

### Transcript-to-complex traits mechanisms of action

Next, we focused on the results of the transcript-to-complex traits analysis to identify examples of transcriptome changes that mediate their phenotypic effect through the proteome. As a first example, we focused on *MANBA* (Figure 7A). After establishing that variants decreasing the gene’s expression colocalized with risk variants for chronic kidney disease [65], the deleterious impact of decreased *MANBA* expression on renal health was recently confirmed in both humans with common expression-altering or rare loss-of-function variants, as well as *Manba* knockout mice [66]. Accordingly, we found that increased *MANBA* transcript had a beneficial impact on kidney damage biomarkers, as it decreased serum urea (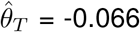, P = 1.2e-5) and cystatin C (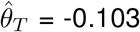, P = 1.6e-12), while increasing estimated glomerular filtration rate (eGFR; 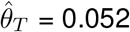, P = 1.4e-6). Importantly, our data shows that these effects are mediated 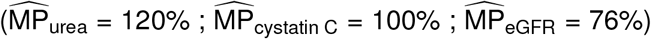 through increased MANBA protein levels (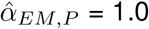, P = 5.2e-10), which in turn affected the aforementioned traits (serum urea: 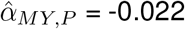, P = 3.2e-5; cystatin C: 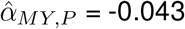, P = 1.9e-9; eGFR: 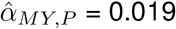, P = 6.3e-9). Furthermore, transcript levels of 3 pseudogenes overlapping the first intron of *MANBA* (RP11-10L12.1 (ENSG00000251288), KRT8P46 (ENSG00000248971); LRRC37A15P (ENSG00000230069)), as well as levels of the adjacent *UBE2D3* antisense RNA RP11-10L12.4 (ENSG00000246560), mediated their phenotypic impact on alkaline phosphatase, cystatin C, diastolic blood pressure, eGFR, and serum urea through decreased MANBA protein levels (Table S7). Overall, this suggests a complex gene-to-phenotype regulation of *MANBA* influenced by nearby non-coding elements.

**Figure 7:**
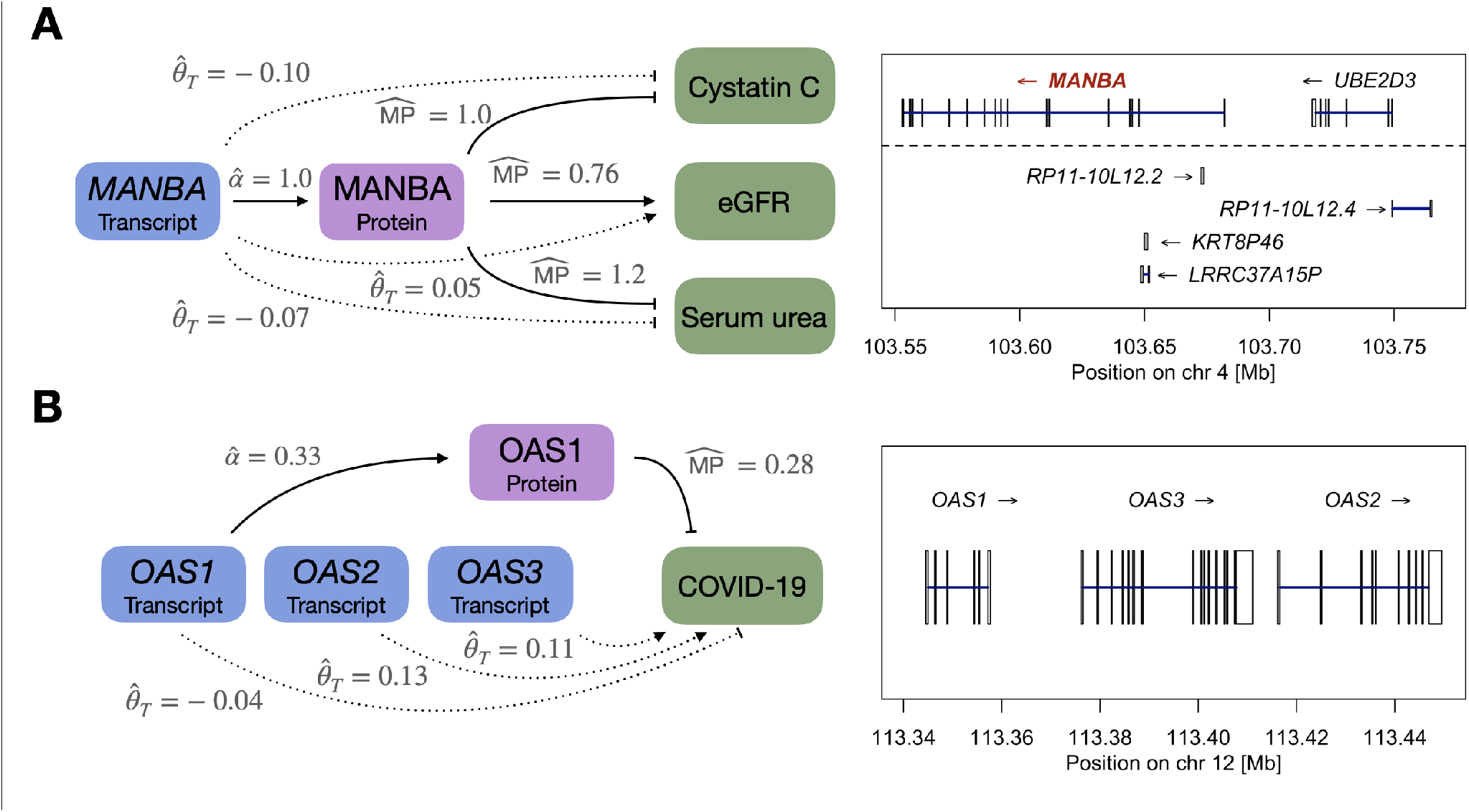
Plausible transcript-protein-trait regulatory mechanisms. **A)** Left: Impact of differential *MANBA* expression on kidney biomarkers through the regulation of its encoded protein. Annotated are the total effect 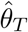 of the *MANBA* transcript levels on the respective outcomes, as well as 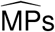 through the encoded protein. Right: Zoom on the *MANBA* region; transcripts below the dashed lines are non-coding and putative negative regulators of MANBA protein levels. **B)** Scheme and locus zoom of the effect of *OAS1/OAS2/OAS3* transcript levels on severe COVID-19 disease. Mediation through the encoded protein could only be tested for *OAS1*.

In contrast, non-coding elements were also found to exert their phenotypic effects through distantly encoded proteins, as illustrated by the transcript originating from the U6 small nuclear RNA 516 pseudogene ENSG00000223313 on chromosome 15, which decreased insulin-like growth factor 1 levels (IGF-1; 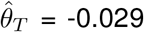, P = 4.0e-7) by decreasing the protein levels of IGF binding protein 3 (IGFBP3; 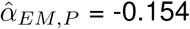, P = 7.6e-4), a well-known regulator of IGF-1’s bioavailability and half-life [67] encoded on chromosome 7 (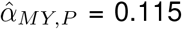, P = 4.0e-7). Alternatively, we observed several cases of protein-coding transcripts affecting traits through proteins in *trans*. For instance, transcript levels of *SUOX*, encoding for a mitochondrial sulfite oxidase, increased lymphocyte count (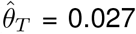, P = 4.7e-5) by positively affecting tyrosylprotein sulfotransferase 2 protein levels (TPST2; 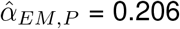, P = 4.0e-4). In turn, TPST2 increased lymphocyte count (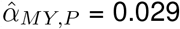, P = 0.02). Both *SUOX* and *TPST2* belong to the KEGG sulfur metabolism pathway (hsa00920). Sulfite oxidase catalyzes the oxidation of sulfite to sulfate [68]. In contrast, sulfate is used by PAPSS1/PAPSS2 to generate 3’-phosphoadenosine-5’-phosphosulfate (PAPS) [69], the main cosubstrate of the sulfotransferase reactions catalyzed by TPST2 [70]. Sulfation of chemokine receptors, which play a critical role in immune function and are widely expressed on lymphocytes, can modulate a receptor’s affinity and/or selectivity for cognate chemokines, as well as mediate pathogen entry [71], establishing the importance of sulfur metabolism for lymphocyte function. Another immune-related example involves the recently established link between the interferon-induced antiviral OAS gene cluster (*OAS1, OAS2, OAS3*) and severe COVID-19 [62, 63]. In line with reports highlighting the protective effect of a Neandertal haplotype associating with increased OAS1 [72, 73], we found that the protective effect against COVID-19 of increased *OAS1* transcript levels (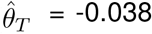, P = 6.9e-8) was mediated 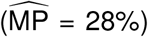 by increased levels of the encoded protein (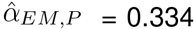, P = 2.0e-23) (Figure 7B). Of note, while protein levels were only available for OAS1, our MR analysis indicated that adjacent and related transcripts *OAS3* (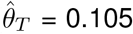, P = 6.8e-8) and *OAS2* (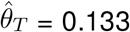, P = 5.2e-3) exerted opposite effects on COVID-19 severity. The opposite effect of *OAS1* and *OAS3* on the outcome reflect previous findings [73] and highlight the complex role of the locus in mediating immunity. Further putative regulatory mechanisms of transcript-to-complex traits through protein levels are shown in Table S10 and were selected based on 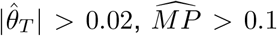 and P_MY,k_ < 0.05. Taken together, these examples illustrate how both protein-coding and non-coding transcripts can exert phenotypic changes through modulation of encoded, as well as *trans* protein levels, suggesting new biological mechanisms.

## Discussion

We presented a framework to quantify mediation of complex trait-impacting effects through multiple omics layers, unravelling nuanced patterns in gene and protein expression regulation. First, we assessed the extent to which DNAm-to-trait effects were mediated by *cis*-transcripts and compared this proportion to the mediation through *cis*-proteins. Evaluating 50 complex traits, the overall adjusted 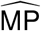 (*i*.*e*.including DNAm-trait pairs with testable mediators in *cis* not under DNAm regulation) through *cis*-transcripts and *cis*-proteins was estimated to be 28.3% and 1.2%, respectively. Simulation studies indicated that the lower sample size of the pQTL dataset (N_pQTL_ ≈ 3,300 vs N_eQTL_ ≈ 30,000) was estimated to result in a relative decrease of 20% in MP, in line with the fact that exposures/mediators with more precise genetic effect estimates are prioritized by MVMR regression models [43]. Despite the fact that ≈6.8x lower number of proteins present in the pQTL dataset (*i*.*e*. fewer testable indirect pathways) than transcripts in the eQTL data, it was not the main reason for the striking difference in the MPs. We demonstrated this by repeating the analysis on a common set of 2,145 transcripts and their encoded proteins, where the adjusted MP through proteins was still ≈10x lower than through transcripts (8.15% vs 0.85%). We suspect that this difference was mainly due to the fact that, on average, proteins were four times less likely to be causally linked to the investigated DNAm site than transcripts, suggesting a tighter link between DNAm and transcript expression than between DNAm and protein levels. This implies a moderate similarity between eQTLs and pQTLs which we confirmed when testing for causal effects between transcript and encoded protein levels: The fraction of testable transcripts linked to their respective protein (when available) at a nominal significance threshold was found to be only 22%. While some of the missing links might be due to the lack of statistical power, it indicates that the transcript to protein regulation is more nuanced than the central dogma of biology would imply, whereby a straight-forward translation from transcripts to proteins by ribosomes is assumed. As a consequence of these weak transcript-to-protein effects, the mediation of transcript-to-trait effects through the encoded protein yielded relatively low MPs (mean = 5.1%). Previous studies reported discrepancies in transcript and protein abundances with explained variances of protein levels by transcript levels ranging from 40 to 85% [74, 75], as well as in eQTL and pQTL co-analyses where only 12 to 40% of the signals were found to be shared [19, 21]. Mechanisms explaining why protein abundance cannot be entirely predicted from transcript levels include protein synthesis delay, transport, degradation, post-transcriptional changes, but also technical variation attributable to measurement instruments [74, 75].

Noteworthily, MR analyses provide directions of estimated causal effects, and two, rather counter-intuitive, observations were made: i) 46.6% of significant DNAm-to-transcript effects were of positive sign (*i*.*e*. DNAm increases transcription) and ii) 20% of significant transcript-to-protein effects were of negative sign (*i*.*e*. high transcript levels decrease protein levels). The first observation is in line with previous genome-wide methylation and gene expression association studies which reported high fractions of positive correlations (30-35%) [48, 46]. While poorly understood [47], several mechanisms have been proposed to explain the phenomenon: preferential binding of some transcription factors to methylated DNA [76, 77], prevention of repressor binding indirectly leading to increased expression through looping DNA [78, 35], or DNAm in the gene body provoking elongation efficiency and preventing spurious initiation of transcription [79]. As to the negative transcript-to-protein effects, which were consistent in the direction when computed with either the INTERVAL or SCALLOP pQTL datasets, literature is more sparse. While negatively correlating gene products have been reported previously [80, 81], this has, to the best of our knowledge, not yet been studied in the context of QTL analyses and remains the topic of future investigations. Finally, MP estimates indicate that DNAm sites typically regulate multiple transcripts in *cis*. Average MPs of 37% suggest that phenotypic DNAm effects are largely mediated through pathways other than local gene expression regulation, especially when the DNAm site is located further away from the TSS of the main transcript mediator. Collectively, these results describe a more diverse picture of the transcription and translation machinery, challenging the classical views of DNAm solely reducing gene expression, and this in the TF region, as well as mRNA levels being a good proxy for protein abundance.

Mapping genetic variants identified in GWAS analyses to biological processes is notoriously difficult [2]. However, systems genetics approaches that integrate multiple omics datasets as a way of leveraging GWAS summary data have proven successful in providing a more complete picture of the path from genotype to phenotype [82]. Here, we demonstrated that our multi-omics framework was able to attribute GWAS signals to biological pathways in loci harbouring multiple genes (e.g. *PARK7* -IBD and *FCERG1*-asthma). A challenge in identifying causal chains through omics layers is the attenuation in the genetic association strengths when moving up along layers. In a linear model, the genetic effect on the phenotype is assumed to be the product of causal effects between the preceding layers and it was previously shown that the variance explained by the top associated QTL of the first layer decreases with each successive omics layer [35]. In line with this observation, the biological examples depicted in Figure 6 visualize the decrease in the genetic associations from the DNAm to the complex trait level. Importantly, integration of both eQTL and pQTL data represent orthogonal approaches in corroborating mediators of DNAm-to-trait or transcript-to-trait effects. Current pQTL datasets lack the sample size and number of proteins to systematically validate regulatory mechanisms found through eQTL integration (e.g. *OAS1*/*OAS2*/*OAS3*-COVID-19). In the future, we expect larger datasets to become available and here presented a proof of concept of how protein-level data can either support mechanistic findings resulting from transcript data or warrant future investigations leading to the discovery of potential new mechanisms of action, implicating other genes.

Throughout the manuscript, we highlighted multiple putative molecular mechanisms of action supported by high MPs through intermediate omics layer and strong literature evidence. More examples can be found in Tables S8-10, including some for which the putative mechanism of action remain strongly debated. For instance, our analyses implicated a DNAm site (cg15133208: chr4:90’757’351) in the TSS region of *SNCA* in Parkinson’s disease (PD) (Figure S23). Many studies have investigated mechanisms involving DNAm, *SNCA* and PD, resulting in conflicting results as to the effect directions. Our results suggest a protective effect of that DNAm site on PD. While supported by studies in the field [83], the assumed DNAm effect on *SNCA* expression is different from our estimated MR effect. Both *SNCA* transcript and SNCA protein levels were estimated to be upregulated in the hypermethylated DNA state, with high *SNCA* levels calculated to decrease PD risk. It is generally assumed that increased *SNCA* expression contribute to PD pathogenesis [84], although blood and brain-specific *SNCA* expression pattern, as well as different isoforms, have been reported to correlate differently with PD [85]. A recent study showed positive correlations between *SNCA* levels and both PD and the related synucleinopathy of Lewy body dementia (LBD) in the temporal cortex, but negative and non-significant ones for LBD and PD in blood, respectively [85]. Another recent GWAS with integrative brain eQTL follow-up analyses indicated that high levels of *SNCA-AS1*, which regulates *SNCA* expression levels, might be protective against LBD [86], suggesting complex regulatory mechanisms governing the locus. Similarly, mechanisms involving proteins in *trans* mediating transcripts-to-trait effects were less straightforward to interpret. Several examples involved non-coding RNA for which functional information is sparse, complicating literature validation.

While our method highlights candidate pathways, several limitations have to be considered. First, like all MR-inverse variance weighting (IVW) analyses, our MR analyses assumed all genetic variants to be valid IVs. We applied Steiger filtering to mitigate the inclusion of pleiotropic IVs that violate independence of the outcome conditional on the exposure and mediators, as well as independence of the mediators conditional on the exposure in the case of variants associated with both the exposure and mediators (third MR assumption; Methods [37]). However, the presence of invalid IVs cannot be excluded and could therefore compromise causal effect estimates [40, 87]. In particular, since selected MR IVs are all in *cis* of the investigated molecular trait, they might be based on a single (pleiotropic) haplotype signal. Conversely, one might argue that the Steiger filter is too stringent if the reverse effect from the mediator on the exposure is biologically unlikely, so that it excludes IVs potentially important in accurately estimating causal effect sizes. Second, we select mediators based on their association to the exposure without taking into account their mediator potential, *i*.*e*. whether or not the mediator is additionally causally linked to the trait. Phrased differently, the selected mediators are simply candidates and such selection serves as a first filter to remove non-mediators. In line with our simulations, it has been shown that extremely large number of such mediator candidates that are not true mediators (92 candidates in total with 88 of them being false mediators) can cause MVMR regression models to fail [43], indicating that our framework is less suitable for large numbers of molecular mediators, unless the selection threshold P_EM_ is made more stringent. Third, our mediation model cannot completely exclude the possibility of reverse effects from the mediator(s) on the exposure. This concern especially applies when considering DNAm as exposure and *cis*-transcripts as mediator(s), since differential transcript levels have been suggested to modulate DNAm levels [35]. We use the largest publicly available mQTL dataset, however, it misses genetic effect sizes of the entire *cis*-region, which would be required to test for reverse or bi-directional effects of transcripts on DNAm. Fourth, with the exception of pQTLs [19], large-scale *trans*-QTL datasets are still lacking, prohibiting genome-wide assessment of mediation and restricting many analyses to *cis*-mediation. Finally, while molecular mechanisms ought to be tissue- or even cell type-specific, QTL data used in this study were all derived from whole blood. It is known that different tissues express different isoforms [88], with many splicing and expression QTLs shown to differ across tissues [89]. Accordingly, MPs for blood biomarkers were generally higher than those for diseases, for which blood might not be the most relevant tissue. Alternatively, this differences might also be due to the fact that indirect pathways, through unmeasured mediators, play a greater role for diseases than for biomarkers. Once tissue-stratified multi-omics datasets of larger sample size become available, more accurate, and potentially higher MPs will be obtained in trait-relevant tissues.

## Conclusion

We quantified the causal connectivity between three omics layers - DNAm, transcript and protein abundance - and their importance in shaping complex traits. We examined regulatory effects of DNAm on gene expression - assessed through both the transcriptome and proteome - which in its complementary use allowed for robust causal inference between molecular and complex traits. Overall, the results indicated that regulatory mechanisms can be more nuanced and complex than suggested by the central dogma of biology, leaving many open questions as to alternative transcription and translation processes. Our integrative omics framework can be extended to other omics-GWAS combinations using the software made available (https://github.com/masadler/smrivw), and provide a powerful tool for mapping GWAS signals to biological pathways and prioritizing functional follow-up experiments.

## Methods

### Univariable and multivariable Mendelian randomization

Univariable Mendelian randomization (MR) was applied to estimate the total causal effect (*θ*_*T*_) and multivariable MR (MVMR) to estimate the direct causal effect (*θ*_*D*_) of an exposure E on an outcome Y. The mediation proportion (MP) was defined as 1 − *θ*_*D*_*/θ*_*T*_. Under the MR assumptions, genetic variants G used as instrumental variables (IVs) must be i) associated with E, ii) independent of any confounder of the E − Y relationship, iii) conditionally independent of Y given E. Independent IVs (*r*^2^ < 0.05) associated with the molecular exposure (P < 1e-6) and located in *cis* (< 1 Mb) allowed the estimation of *θ*_*T*_ using an inverse-variance weighted (IVW) method assuming equal weights given the standardization of the data and accounting for correlated instruments [90]:

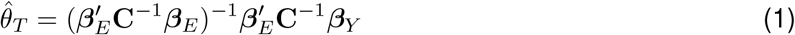

where *β*_*E*_ and *β*_*Y*_ are vectors of genetic effect sizes obtained from summary statistics for E and Y, respectively. C is the linkage disequilibrium (LD) matrix with pairwise correlations between IVs estimated from the UK10K reference panel [91]. Prior to the causal effect calculation, IVs were filtered to fulfill the MR Steiger criterion of no larger Y than E genetic effects [37] and were thus required to pass a threshold 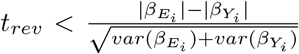 with *t*_*rev*_ set at −2, equivalent to a one sided test p-value threshold of 0.023. IVs not passing this threshold are prone to violating the third MR assumption of horizontal pleiotropy since they are more directly linked to the outcome. As a result MR estimates including such IVs would potentially mix up forward and reverse causal effects. The standard error (SE) of *θ*_*T*_ can be approximated by the Delta method [92]:

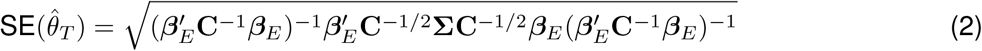

where Σ is a diagonal matrix with each diagonal element *i* equalling the maximum of the regression variance *s*^2^ and 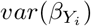 [93].

Through the inclusion of mediators M_k_ and their associated *cis* genetic variants (*r*^2^ < 0.05, P < 1e-6), *θ*_*D*_ can be estimated analogously to *θ*_*T*_ using a multivariable regression model [41] as the first element of *θ*_*D*_:

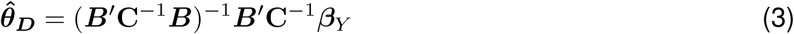

where *B* is a matrix with *k* + 1 columns containing the effect sizes of the IVs on the exposure in the first column and on each mediator in the subsequent columns. The remaining elements of *θ*_*D*_ represent the direct effects of the mediators on the outcome and were referred to as *α*_*MY,k*_. In the estimation of MPs, we were not interested in *α*_*MY,k*_ values *per se*, but we took these effect sizes into account for inferring molecular mechanisms. If the number of mediator-associated instruments was sufficient (≥ 3) to conduct a univariable MR from the mediator on the outcome, we estimated *α*_*MY,k*_ from this analysis instead, since computed on a single regressor, narrower CIs are obtained.

This MVMR model does not allow for the presence of a causal effect from the mediators on the outcome via the exposure, and we therefore conducted several Steiger filtering steps on the IVs. In addition to meeting the Steiger criterion described above, exposure-associated IVs were required to pass that same threshold *t*_*rev*_ of no larger mediator than exposure effects for each of the mediators M_k_. Similarly, to mitigate reverse causal effects from the outcome on the mediators, mediator-associated instruments with larger Y than M effects were removed if not passing the *t*_*rev*_ threshold. The SE of 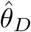 was derived analogously to the univariable form as shown in [29].

### Omics and trait summary statistics

We used mQTL data from the GoDMC consortium (N = 32,851) [16], which contains > 170,000 whole blood DNAm sites with at least one significant *cis*-mQTL (P < 1e-6, < 1 Mb from the DNAm site, N > 5,000). *Cis*-eQTL data were taken from the eQTLGen consortium (N = 31,684) [18] which includes *cis*-eQTLs (< 1 Mb from gene center, 2-cohort filter) for 19,250 transcripts (16,934 with at least one significant *cis*-eQTL at FDR < 0.05 corresponding to P < 1.8e-05). *Cis*- and *trans*-pQTL data were from the INTERVAL study (N = 3,301) [19]. SomaLogic aptamers (N_SOMAmers_ = 3,283) quantified the levels of 2,977 proteins and complexes with a UniProt ID. After removing protein complexes (N_SOMAmers_ = 42), sex chromosome encoded proteins (N_SOMAmers_ = 113), and UniProt IDs that could not be mapped to an EnsemblID (N_SOMAmers_ = 6), 2,838 proteins remained of which 696 had at least one significant *cis*-pQTL (P < 1e-6, < 1 Mb from the protein-encoding gene center, N > 2,000). If two SOMAmers mapped to the same protein, the one with the strongest transcript-to-protein causal MR effect was retained (see omics-to-omics MR analysis). If the transcript was not available in the eQTLGen dataset or did not have any significant IVs, the SOMAmer with the highest number of significant *cis*-pQTLs (or *trans*-pQTLs if no *cis*-pQTLs were present) was chosen. Mapping from UniProt to Ensembl identifiers was done through the UniProt REST API [94] and genomic coordinates were retrieved from the Ensembl REST API (GRCh37 build) [95]. Exact mapping of SOMAmer-UniProt-Ensembl identifiers is provided in Table S5. A total of 2,145 transcript-encoded protein pairs were present in both the eQTL and pQTL datasets.

GWAS summary statistics for outcome traits came from the largest (N_average_ > 320,000), predominantly European-descent, publicly available studies, as listed in Table S1.

Prior to each mediation analysis, exposure and mediator omics, GWAS and the reference panel data were harmonized. The analysis was conducted on autosomal chromosomes, and palindromic single nucleotide variants (SNPs), as well as SNPs with an allele frequency difference > 0.05 between any pairs of datasets were removed. If allele frequencies were not reported by the GWAS summary statistics, allele frequencies from the UK Biobank were used. Z-scores of summary statistics (molecular and outcome GWAS) were standardized by the square root of the sample size to be on the same SD scale.

### DNAm-to-trait mediation analysis

First, univariable MRs were conducted to estimate the total causal effect 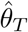 of the DNAm sites on each trait, assessing ∼50,000 DNAm probes with ≥ 5 independent mQTLs after harmonization of the datasets (*r*^2^ < 0.05). DNAm probes significantly associated to the outcome (P_T_ < 0.05/50000 = 1e-6) were clumped based on the p-value of the total causal effect 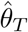, P_T_ (distance-pruning at 1 Mb), to be independent of each other.

Second, MVMR analyses were performed to estimate the direct effect 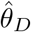. Potential transcript mediators in *cis* of the DNAm exposure probe (± 500kb) were extracted and causal effects *α*_*EM,k*_ of the DNAm probe on these transcripts were assessed by univariable MR. Transcripts satisfying P_EM,k_ < P_EM_ (default P_EM_ = 0.01, with 0.05 and 1e-3 being tested as well) were included as mediators, as well as their associated SNPs as additional instruments. Steiger filtering was applied as described previously and IVs were clumped based on a rank score determined as follows: 1) for each mediator, IVs were ranked according to their association p-value to the mediator and assigned an integer score, 2) for each IV, a final score was calculated as the sum of its individual mediator scores. Following the establishment of the *B* effect size matrix, 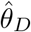 was calculated, as well as 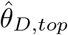 which was estimated from a MVMR model that includes a single mediator, namely the transcript with the lowest P_EM,k_. If no transcript causally associated to the DNAm probe, mediation is not detectable, hence 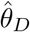 was set to 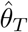 for that probe (inclusion of such probes in MP calculation was termed “adjusted mediation proportion”). As the Steiger filter removed exposure-associated instruments with larger mediator than exposure effects (see “Univariable and multivariable Mendelian randomization”), the number of initial exposure-associated instruments (*m*_*E*_ ≥ 5) could decrease. Therefore, to avoid scenarios of reverse causality where the mediator exerts an effect on the outcome through the exposure, we required ≥ 3 exposure-associated IVs.

We additionally conducted mediation analyses on independent mediators. To this end, selected mediators (those that passed P_EM_) were clumped at various correlation thresholds R_med_ (default R_med_ < 0.3, with 0.2 and 0.1 being tested as well). Correlations among the mediators were calculated based on QTL effect sizes of independent exposure and mediators IVs and priority was given to the mediator with the lowest P_EM,k_.

The mediation proportion (MP) was calculated by regressing 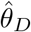 on 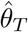 to estimate for the unmediated proportion, 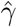, which after correcting for regression dilution bias (Equation 4):

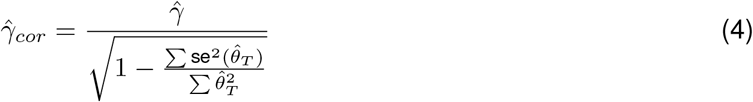

yielded 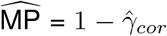 for a defined set of DNAm-trait pairs. MVMR analyses were repeated on the selected DNAm-trait pairs through proteins in *cis* following the same mediator and IV filtering steps as described above.

### Transcript-to-trait mediation analysis

MPs for transcript-to-trait mediation analyses were calculated similarly to DNAm-to-trait MPs. Briefly, we first computed total causal effects 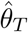 of transcripts on traits for ∼ 11,000 transcripts with ≥ 5 independent (*r*^2^ < 0.05) and significant eQTLs (P < 1e-6), ∼ 1,200 of which had an encoded protein in the pQTL dataset. For each trait, significant transcripts (P_T_ < 0.05/1,000 = 5e-5) were selected. Second, MVMR analyses were conducted, where for each transcript, mediators were defined as i) the encoded protein or ii) the encoded protein plus any other protein in *trans* among a set of 696 proteins with ≥ 1 significant (P < 1e-6) pQTL that satisfied P_EM,k_ < P_EM_ (default P_EM_ = 1e-3, with 1e-2 and 1e-4 being tested as well). If more than 10 proteins satisfied the condition, the ten most strongly associated were retained. Associated pQTLs were included as IVs and following Steiger filtering, instruments were pruned as described in the DNAm-to-trait mediation analysis section. Effect sizes of mediator-associated IVs that were not significant (P > 1e-6) for a given mediator were shrunk to 0 [96]. Direct effects 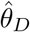 were calculated using encoded proteins (if available) as mediators in addition to selected *trans* proteins. Additionally, direct effects were calculated using only encoded proteins as mediators. Finally, 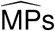 were calculated by aggregating all transcript-trait pairs as specified in each sub-analysis, and regressing 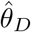 on 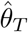 while accounting for regression dilution bias (Equation 4).

### Omics-to-omics MR analysis

MR causal effects between two molecular traits were calculated following the same procedure than in the univariable MR to calculate total effects 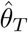. First, independent (*r*^2^ < 0.05) and significant (P < 1e-6) exposure IVs were selected and IVs not passing the aforementioned Steiger filter were discarded. MR causal effects were then computed based on Equation 1.

#### DNAm-to-transcript MR analysis

MR effects between DNAm sites and transcripts in *cis* (± 500kb) with ≥ 3 exposure IVs were calculated. Pearson correlation coefficient with previously reported DNAm-transcript correlations [48] was calculated on common DNAm-transcript pairs. DNAm probe annotations with respect to the assessed transcript were from the IlluminaHumanMethylation450kanno.ilmn12.hg19 R package [97].

#### Transcript-to-encoded-protein MR analysis

On the common transcript-encoded protein pairs, causal effects were calculated for transcripts with ≥ 3 independent eQTLs (*r*^2^ < 0.05). When comparing causal effects obtained from the INTERVAL and SCALLOP pQTL dataset, we additionally included transcripts with a single eQTL.

### Simulation studies

We conducted simulation studies to assess the robustness of our model and to identify sources of bias in the estimated MP. Two simulation settings were set up: one replicating the DNAm-to-trait via transcripts in *cis* mediation analysis and one replicating the transcript-to-trait via proteins in *trans* mediation analysis. Both scenarios were simulated under the same model, but with different parameter settings (Figure S1, Table S2).

We considered an exposure with heritability 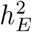 and *m*_*E*_ independent IVs. Effect sizes 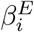 for *m*_*E*_ IVs were drawn from a normal distribution 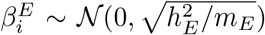 and rescaled to total 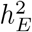. *N*_*med*_ potential mediators were simulated, among which *N*_*med,sig*_ were contributing to the indirect effect *θ*_*M*_. Each mediator *k* associated with *m*_*M*_ IVs with direct effects 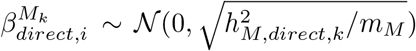 rescaled to 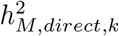, the direct heritability of the mediator that does not take into account the additional heritability coming through the exposure. Direct heritabilities were sampled from a uniform distribution 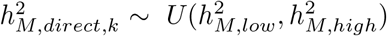. Causal effects of the exposure on the mediator (*α*_*EM,k*_) and of the mediator on the outcome (*α*_*MY,k*_) for *N*_*med,sig*_ mediators were drawn from a bivariate normal distribution *α*_*EM,k*_, *α*_*MY,k*_ ∼ 𝒩(0, Σ) with Σ the covariance matrix:

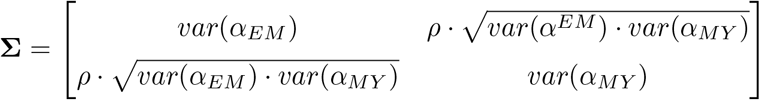

where *ρ* is the correlation between *α*_*EM,k*_ and *α*_*MY,k*_. For the remaining *N*_*med*_ - *N*_*med,sig*_ mediators, *α*_*EM,k*_ and *α*_*MY,k*_ causal effects were set to zero. The vector of effect sizes 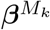 of size *m*_*E*_ + *N*_*med*_ · *m*_*M*_ for each mediator *k* was constructed to have effect sizes equalling 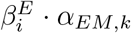 for *m*_*E*_ exposure SNPs and effect sizes equalling 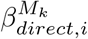 for *m*_*M*_ mediator-associated SNPs. The effect sizes of remaining IVs associated to mediators *i* ≠ *k* were set to zero. Likewise, effect sizes of the *N*_*med*_ · *m*_*M*_ IVs on the exposure in the *β*^*E*^ vector were set to zero.

The indirect effect *θ*_*M*_, direct effect *θ*_*D*_ and total effect *θ*_*T*_ were calculated as:

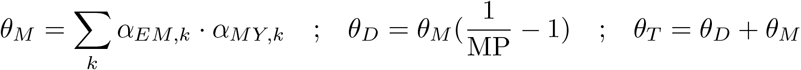

These quantities allowed to design the outcome effect size vector *β*^*Y*^:

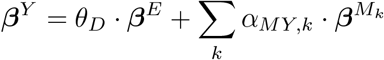

For each scenario, we simulated 300 data sets to each time get *β*^*E*^, 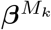 and *β*^*Y*^. Normally distributed noise, as a function of the sample size N, 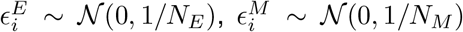 and 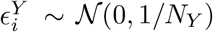 was added to each simulated vector. To approximate our real data, exposure effect sizes of SNPs serving as mediator instruments were set to zero again. We then estimated for each model 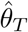 and 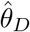 by including mediators that satisfied P_EM_ (p-value of the causal effect from the exposure on the mediator). Causal effects 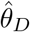 were regressed on 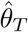 to estimate the coefficient 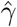 which after accounting for regression dilution (Equation 4) allowed to obtain the estimated 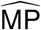.

### Comparing mediation proportions

To test the statistical significance between 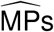 estimated on two different sets of exposure-trait pairs (e.g. 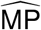 of a given physiological category vs all categories combined) or on the same exposure-trait pairs, but with different parameter settings (e.g. changing P_EM_), we make use of 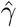 and its corresponding standard error 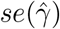 obtained from regressing 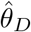 on 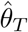 (both of which being corrected for regression dilution (Equation 4)) to yield 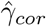 and 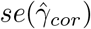. We then perform a two-sided z-test based on the following test statistic:

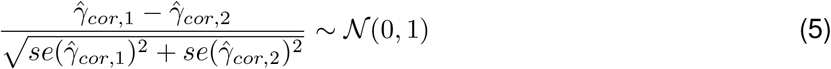

A significant difference between 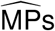 was claimed if the two-sided p-value was below 0.05.

## Availability of data and materials

QTL datasets can be downloaded at the following websites: mQTLs (http://mqtldb.godmc.org.uk/downloads), eQTLs (https://www.eqtlgen.org/cis-eqtls.html), pQTLs (http://www.phpc.cam.ac.uk/ceu/proteins/. The list of GWAS summary statistics used is in Table S1, all of which are all from the public domain.

Software to conduct univariable MR-IVW (molecular trait → outcome, molecular trait 1 → molecular trait 2) and multivariable MR-IVW (molecular trait 1 → molecular trait 2 → outcome) can be found at https://github.com/masadler/smrivw. Source code (C++, released under GPL v3 license) and executable file (for Linux platforms, released under MIT license) are provided which rely on functionalities and the data management architecture of the SMR software (https://cnsgenomics.com/software/smr [35]). The provided documentation hosted on the github repository guides users in reproducing the mediation results and conducting univariable and multivariable MR on their own combinations of QTL and GWAS datasets.

## Competing interests

The authors declare that they have no competing interests.

## Authors’ contributions

M.C.S., E.P. and Z.K. conceived and designed the study. M.C.S. performed statistical analyses. E.P. provided guidance on statistical analyses. Z.K. supervised all statistical analyses. All the authors contributed by providing advice on interpretation of results. C.A. contributed with the biological interpretation of the results. M.C.S., E.P. and Z.K. drafted the manuscript. C.A. contributed to the writing of specific sections. All authors read, approved, and provided feedback on the final manuscript.

## Acknowledgements

LD was calculated based on the UK10K data resource (EGAD00001000740, EGAD00001000741). Computations we performed on the JURA cluster of the University of Lausanne. We would also like to thank Kaido Lepik for critical comments on the manuscript.

## Notes

### Competing Interest Statement

The authors have declared no competing interest.

