## Supplementary Materials for "Quantifying mediation between omics layers and complex traits"

### Supplementary Figures

#### Simulation studies

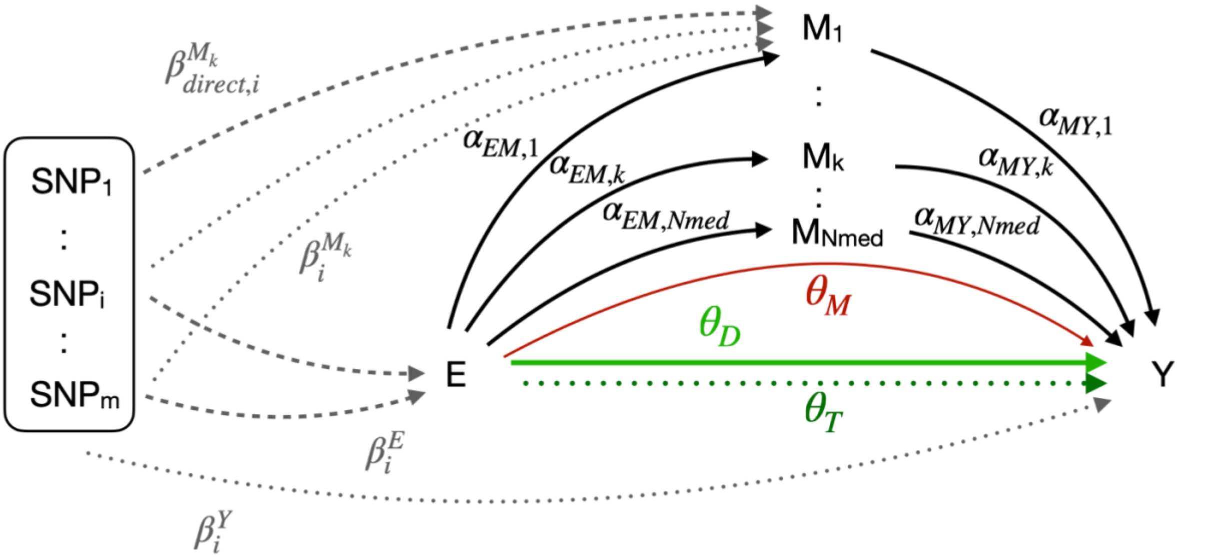

**Supplementary Figure 1**. Model used in the simulation settings to estimate the total, direct and indirect causal effects ($\Theta_T$, $\Theta_D$ and $\Theta_M$, respectively). Genetic variants (SNPs) are either directly associated (dashed arrow) with the exposure E or mediators $M_k$ (1 to $N_{med}$), or indirectly (dotted arrow) with $M_k$ through E. The genetic effect sizes are denoted by β, where $\beta^E$ are direct effects to E, $\beta^{M_k}$ total effects to $M_k$ made of the direct effects $\beta^{M_k}_{direct}$ to $M_k$ and the indirect effects through E, and $\beta^Y$ are total effects to the outcome Y through either E or M. Causal effects from E to $M_k$ are denoted by $\alpha_{EM,k}$ and causal effects from M to Y by $\alpha_{MY,k}$.

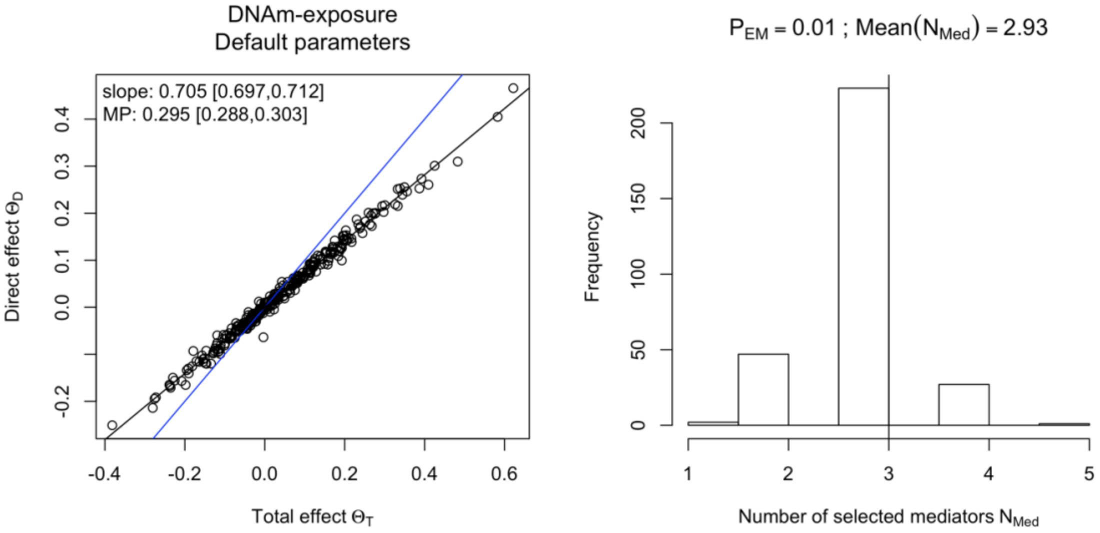

**Supplementary Figure 2.** Simulation results in the DNAm-exposure setting (default settings as indicated in Table S2). 300 exposure-outcome pairs were simulated and for each a direct and total effect was estimated. The slope (corrected for regression dilution) and the mediation proportion together with the 95% CI are displayed in the plot area (blue line represents the identity line). Mediators were selected based on a p-value threshold $P_{EM}$ and the distribution of the number of mediators selected is shown in the histogram. The true number of relevant mediators was 3 and the true MP was 0.3 (Table S2).

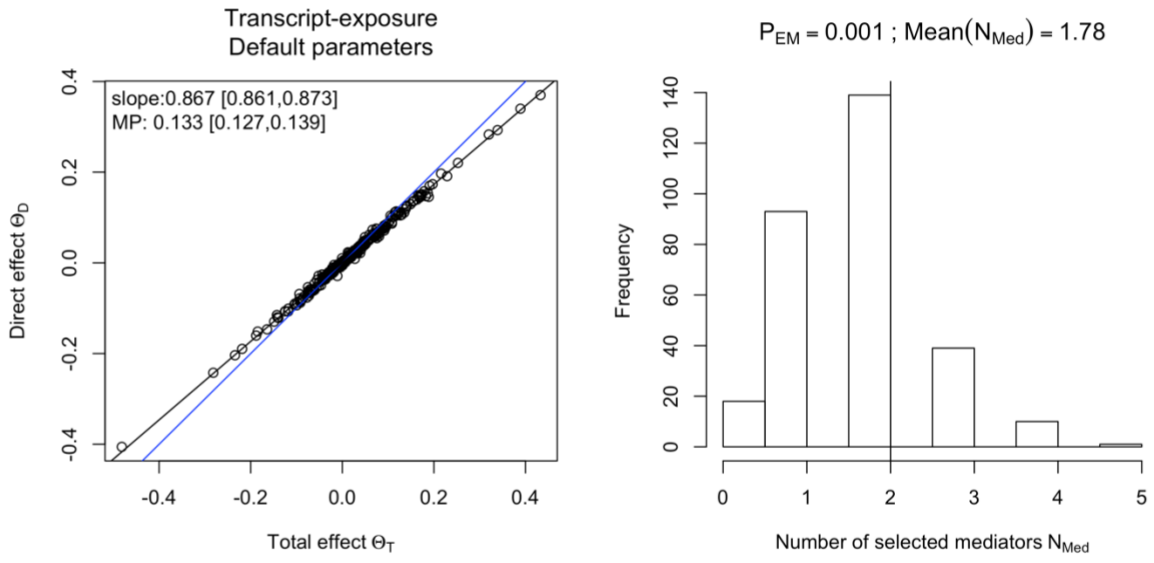

**Supplementary Figure 3.** Simulation results in the transcript-exposure setting (default settings are indicated in Table S2). Again 300 exposure-outcome pairs were simulated and the number of relevant mediators was 2 with the true MP being 0.15. The slope (corrected for regression dilution) and the mediation proportion together with the 95% CI are displayed in the plot area (blue line represents the identity line). Mediators were selected based on a p-value threshold $P_{EM}$ and the distribution of the number of mediators selected is shown in the histogram.

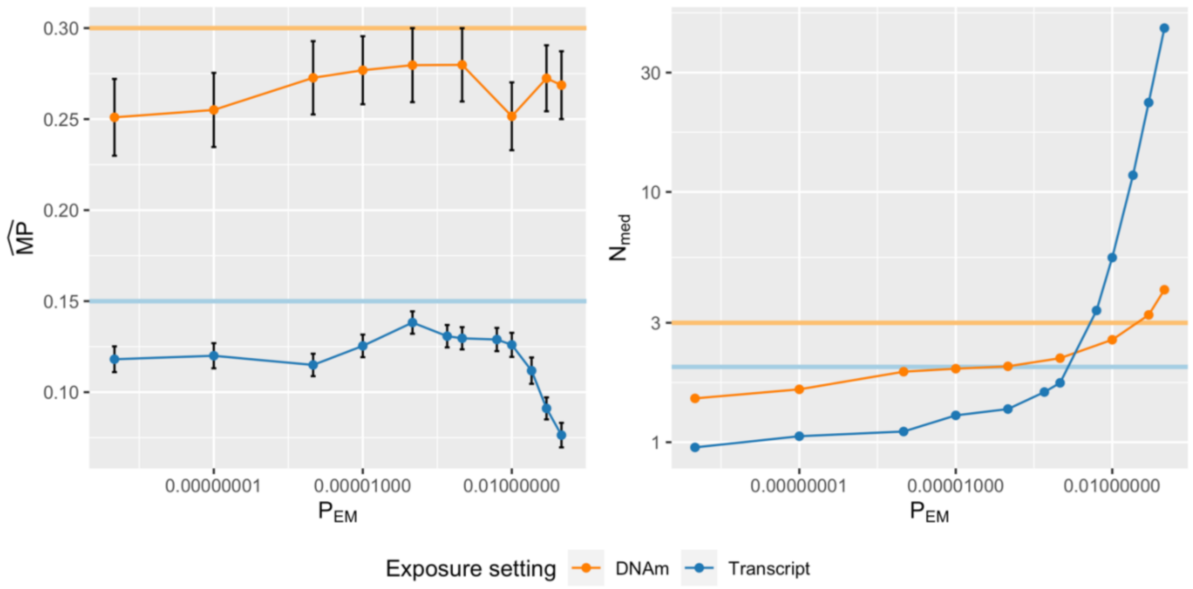

**Supplementary Figure 4**. Simulation results in DNAm- and transcript-exposure settings to assess the impact of the $P_{EM}$ threshold on the A) estimated MP and B) number of selected mediators. For a given mediator sample size, 300 exposure-outcome pairs were simulated on which an MP and 95\% CI (error bars) were estimated. The true MP of the model was 0.3 and 0.15, and the true number of relevant mediators was 3 and 2 in the DNAm-exposure and transcript-exposure setting, respectively, as indicated by the solid horizontal lines.

### DNAm-to-trait through transcripts/proteins in *cis* mediation analysis

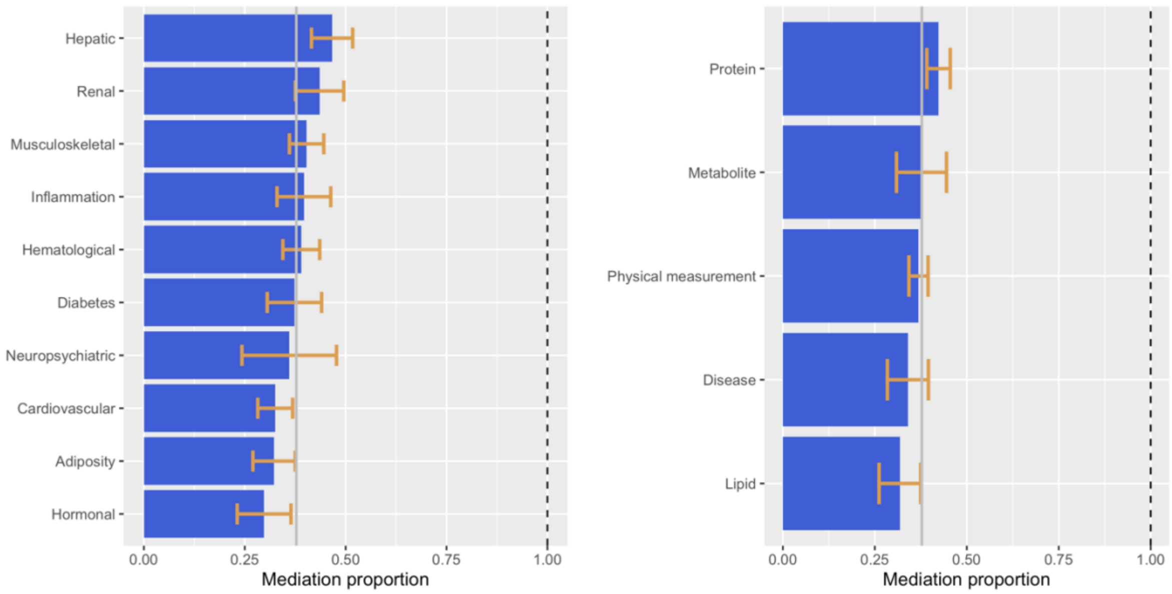

**Supplementary Figure 5.** Mediation proportion of traits grouped by physiological (left) and structural (right) categories in the DNAm-to-trait through transcripts in *cis* mediation analysis (see Table S1 for trait classification). The grey vertical line denotes the mean mediation proportion across all DNAm-trait pairs. 95% confidence intervals are represented by the orange error bars.

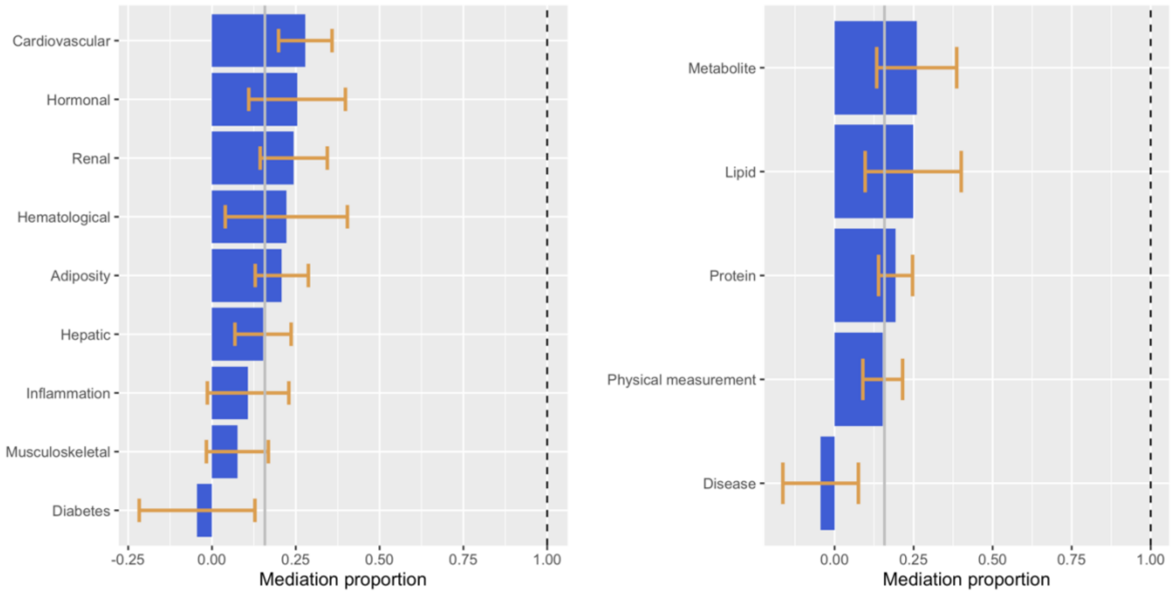

**Supplementary Figure 6.** Mediation proportion of traits grouped by physiological (left) and structural (right) categories in the DNAm-to-trait through proteins in *cis* mediation analysis (see Table S1 for trait classification). Only categories with at least 10 DNAm-trait pairs were evaluated. The grey vertical line denotes the mean mediation proportion across all DNAm-trait pairs. 95% confidence intervals are represented by the orange error bars.

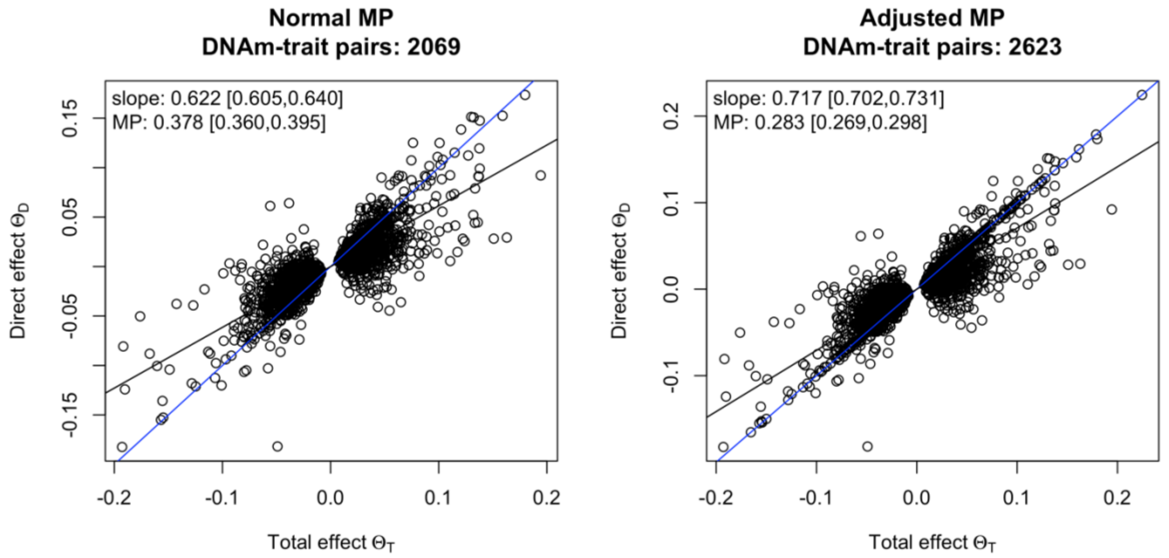

**Supplementary Figure 7.** Comparison of the normal and the adjusted MP in the DNAm-to-trait through transcripts in *cis* mediation analysis. In the calculation of the normal MP, DNAm-trait pairs were evaluated for which there was at least 1 transcript in the *cis* region causally associated to the DNAm site under scrutiny. In the adjusted MP calculation, all DNAm-trait pairs for which there was at least 1 transcript in the *cis* region, but not necessarily causally linked to the DNAm site were evaluated. Here, the direct effects of DNAm-trait pairs for which there was no mediator were set to the total effect. Plotted is the direct effect against the total effect together with the slope adjusted for regression dilution in black and the identity line in blue.

### DNAm-to-trait analysis: Comparison of transcripts and proteins in *cis* as mediators

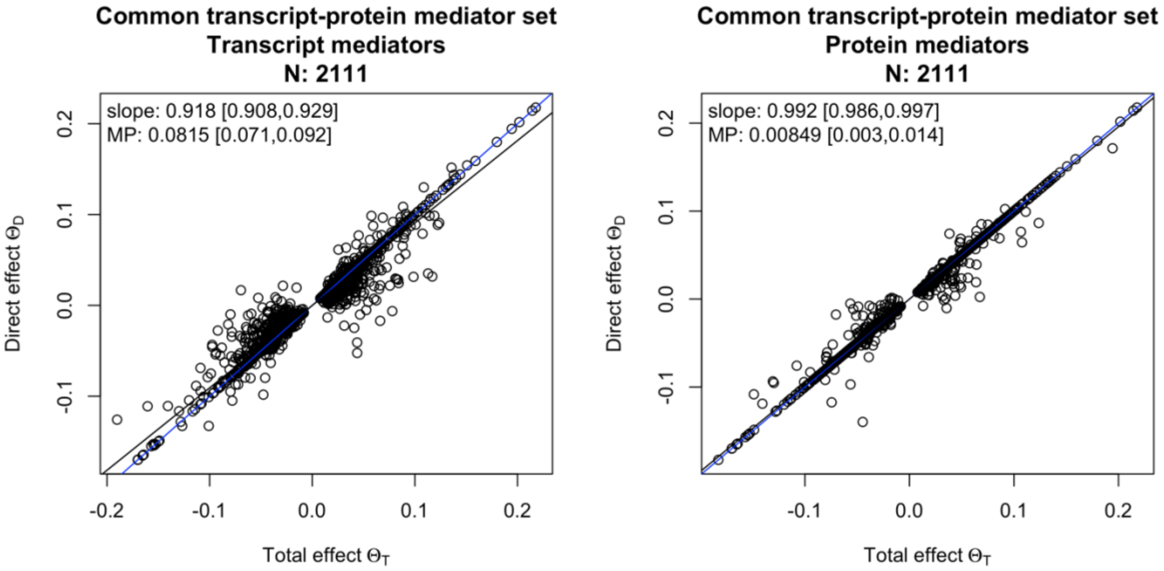

**Supplementary Figure 8.** Calculation of the adjusted mediation proportion on the common transcript-protein mediator set. DNAm-trait pairs (number indicated in the title by N) were evaluated for which there was at least 1 transcript/protein in the *cis* region. Plotted is the direct effect against the total effect together with the slope adjusted for regression dilution in black and the identity line in blue. The direct effects of DNAm-trait pairs for which there was no mediator causally associated to the DNAm site were set to the total effect.

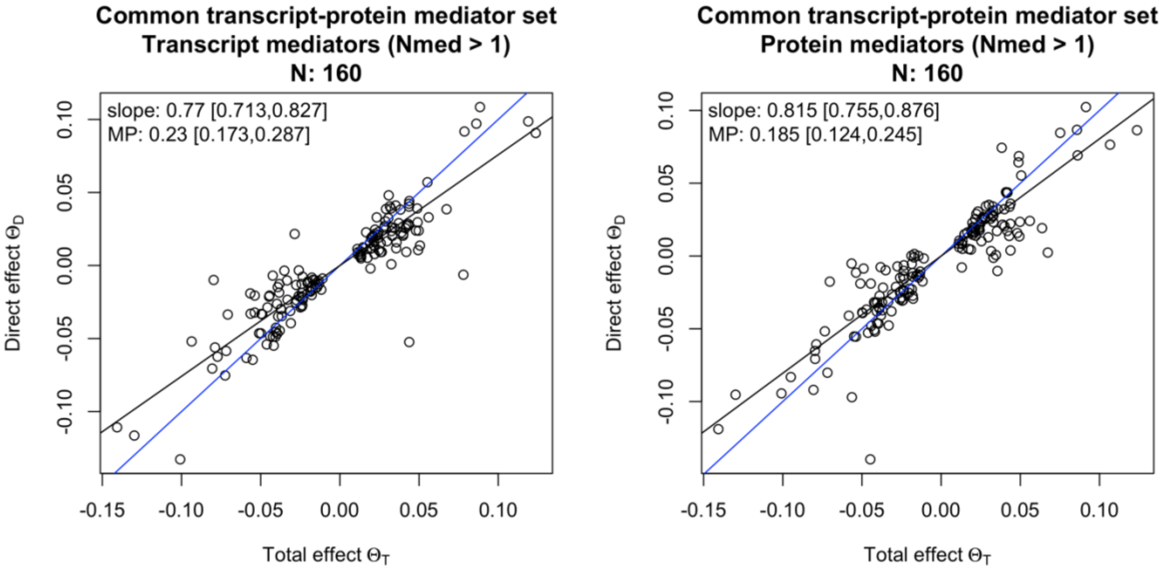

**Supplementary Figure 9.** Calculation of the mediation proportion on the common transcript-protein mediator set. DNAm-trait pairs (number indicated in the title by N) were evaluated for which there was at least 1 transcript and protein in the *cis* region causally associated to the DNAm site. Plotted is the direct effect against the total effect together with the slope adjusted for regression dilution in black and the identity line in blue.

### DNAm-to-trait mediation analysis: Sensitivity analyses

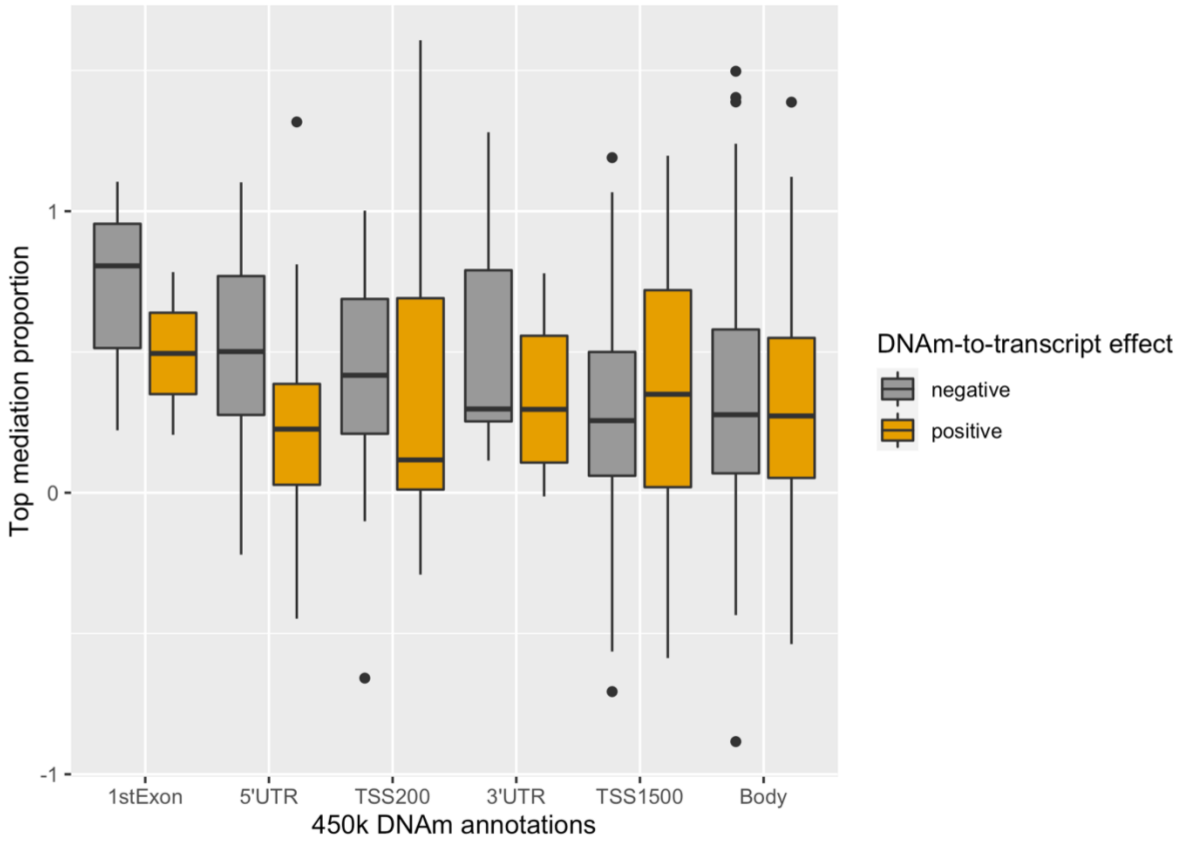

**Supplementary Figure 10.** Boxplots representing the top mediation proportion ($MP_{top}$) stratified by DNAm site location with respect to the top mediator and by the causal effect direction of the DNAm on the transcript level. The annotation groups are shown in decreasing order with respect to the mediation proportion (negative and positive DNAm-to-transcript effect pairs combined).

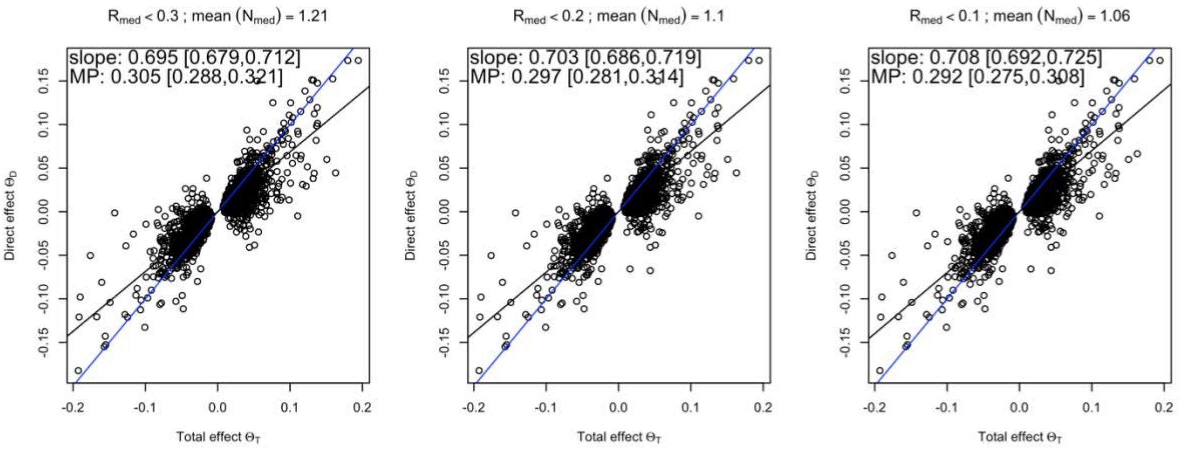

**Supplementary Figure 11**. DNAm-to-trait through transcripts in *cis* mediation analysis with uncorrelated mediators at different $R_{med}$ thresholds (0.3, 0.2, and 0.1 from left to right). $R_{med}$ is the maximum correlation between the mediators for a given exposure-outcome pair. As this threshold decreases, the average number of selected mediators ($N_{med}$) decreases. The slope (corrected for regression dilution) and the mediation proportion together with the 95% CI are displayed in the plot area (blue line represents the identity line).

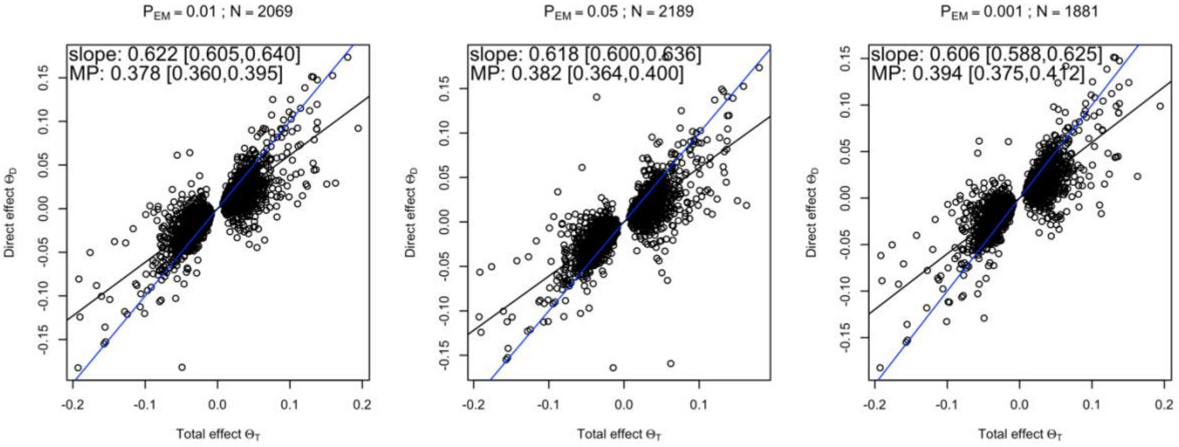

**Supplementary Figure 12.** DNAm-to-trait through transcripts in *cis* mediation analysis with different $P_{EM}$ thresholds to select mediators (0.01, 0.05 and 0.001 from left to right). The calculation of the MP is only done on DNAm-trait pairs (N pairs) with at least 1 transcript in the *cis* region causally associated to the DNAm site. The slope (corrected for regression dilution) and the mediation proportion together with the 95% CI are displayed in the plot area (blue line represents the identity line).

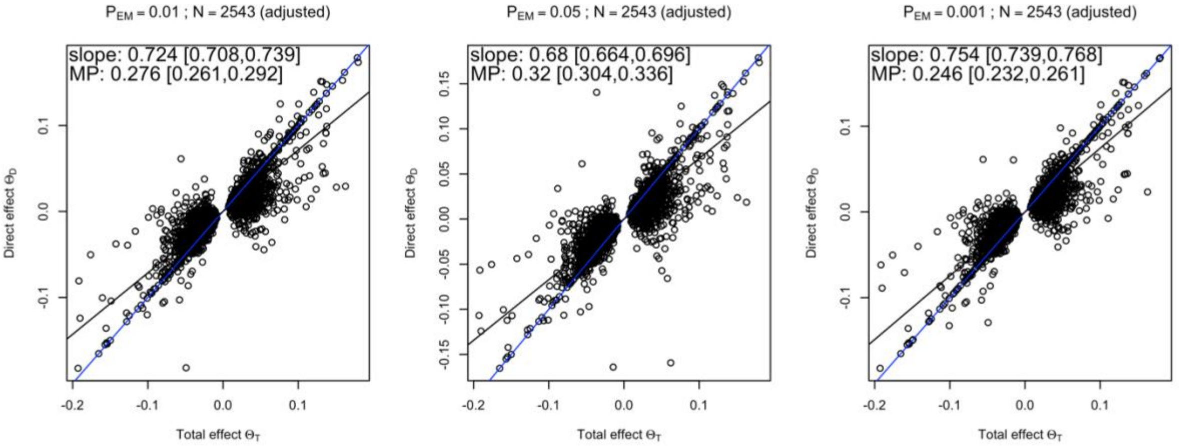

**Supplementary Figure 13.** DNAm-to-trait through transcripts in *cis* mediation analysis with different $P_{EM}$ thresholds to select mediators (0.01, 0.05 and 0.001 from left to right). The adjusted MP is calculated on all DNAm-trait pairs (N pairs) with least 1 transcript in the *cis* region (not necessarily causally associated to the exposure) and for which a mediation analysis could be performed in all three settings (the number of instrumental variables being the limiting factor). The slope (corrected for regression dilution) and the mediation proportion together with the 95% CI are displayed in the plot area (blue line represents the identity line).

#### Transcript-to-trait through proteins mediation analysis

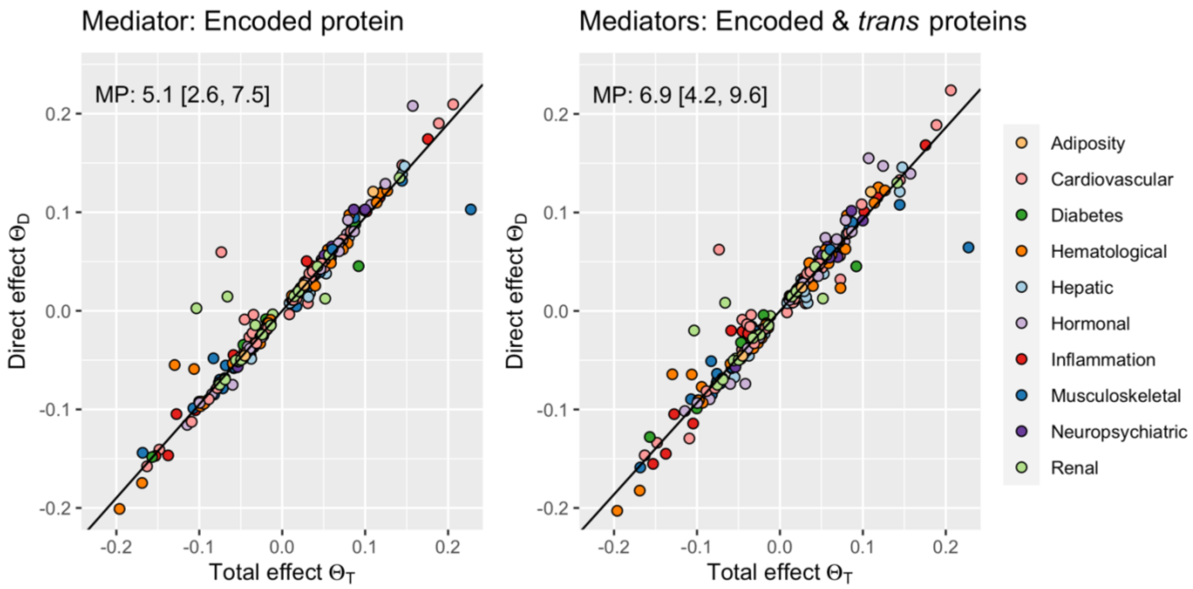

**Supplementary Figure 14.** Transcript-to-trait through proteins mediation analysis. Left: through the encoded protein only. Right: through encoded and any additional *trans* proteins causally linked to the investigated transcript. The slope (corrected for regression dilution) is plotted, and the global MPs in % with 95% CI are indicated in the plotting area. Transcript-trait pairs are colour-coded according to the physiological category of the trait (see Table S1 for trait classification).

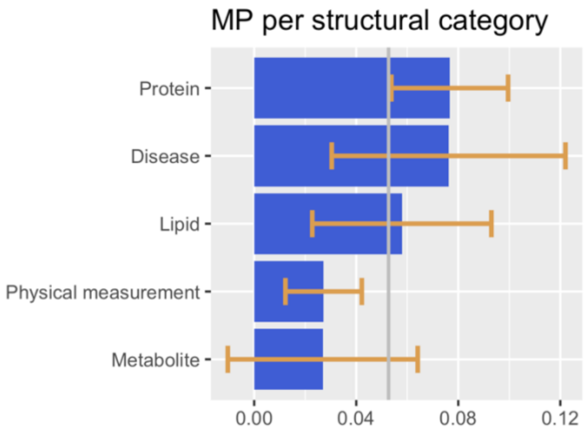

**Supplementary Figure 15.** Mediation proportion of traits grouped by structural category in the transcript-to-trait through proteins (encoded, if present, and *trans*) mediation analysis (see Table S1 for trait classification). The grey vertical line denotes the mean mediation proportion across all transcript-trait pairs. 95% confidence intervals are represented by the orange error bars.

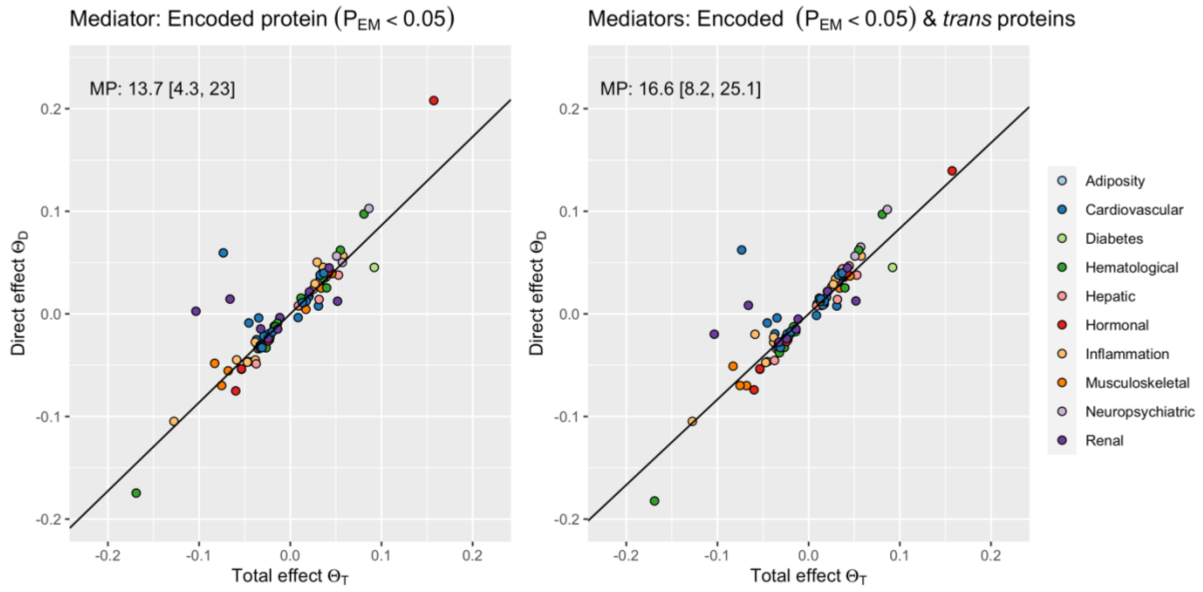

**Supplementary Figure 16.** Transcript-to-trait mediation analysis of transcripts for which the encoded protein was nominally significantly associated to the transcript. The left panel shows the mediation results through the encoded protein alone, and the right panel through the encoded protein plus any additional protein in *trans* causally associated to the transcript. The MP in % calculated on all transcript-trait pairs together with the 95% CI is indicated in the plotting area.

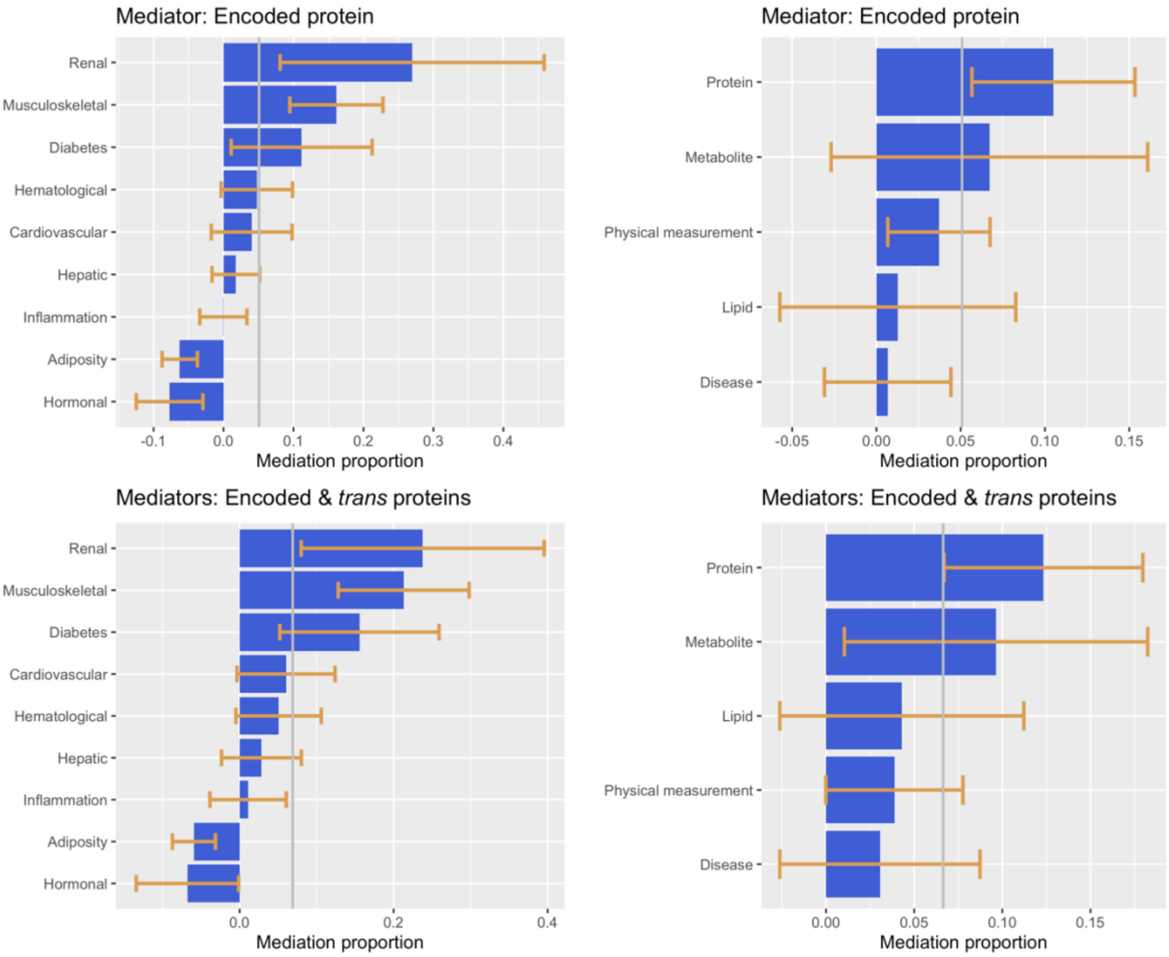

**Supplementary Figure 17.** Mediation proportion of traits grouped by physiological and structural categories in the transcript-to-trait through proteins mediation analysis (see Table S1 for trait classification). The grey vertical line denotes the mean mediation proportion across all the transcript-trait pairs. 95% confidence intervals are represented by the orange error bars. The top row shows the mediation results through the encoded protein alone and the bottom row through the encoded protein plus any additional protein in *trans* causally associated to the transcript. Only categories with at least 10 transcript-trait pairs were evaluated.

### Transcript-to-trait through proteins mediation analysis: Sensitivity analyses

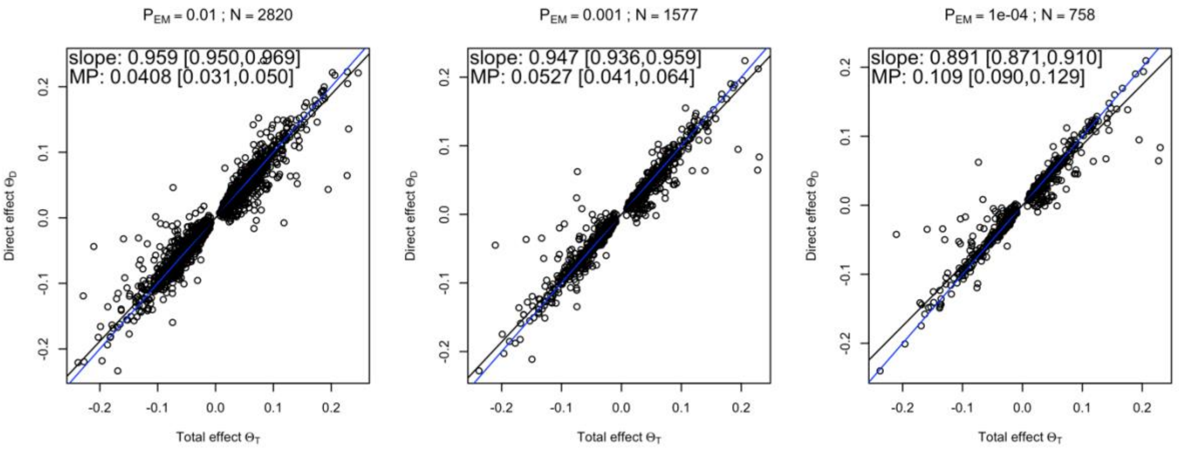

**Supplementary Figure 18.** Transcript-to-trait through proteins (encoded, if present, plus additional *trans* proteins) mediation analysis with different $P_{EM}$ thresholds to select mediators. The calculation of the MP is only done on transcript-trait pairs (N pairs) with at least 1 mediator protein.

### Comparison between the INTERVAL and SCALLOP pQTL datasets

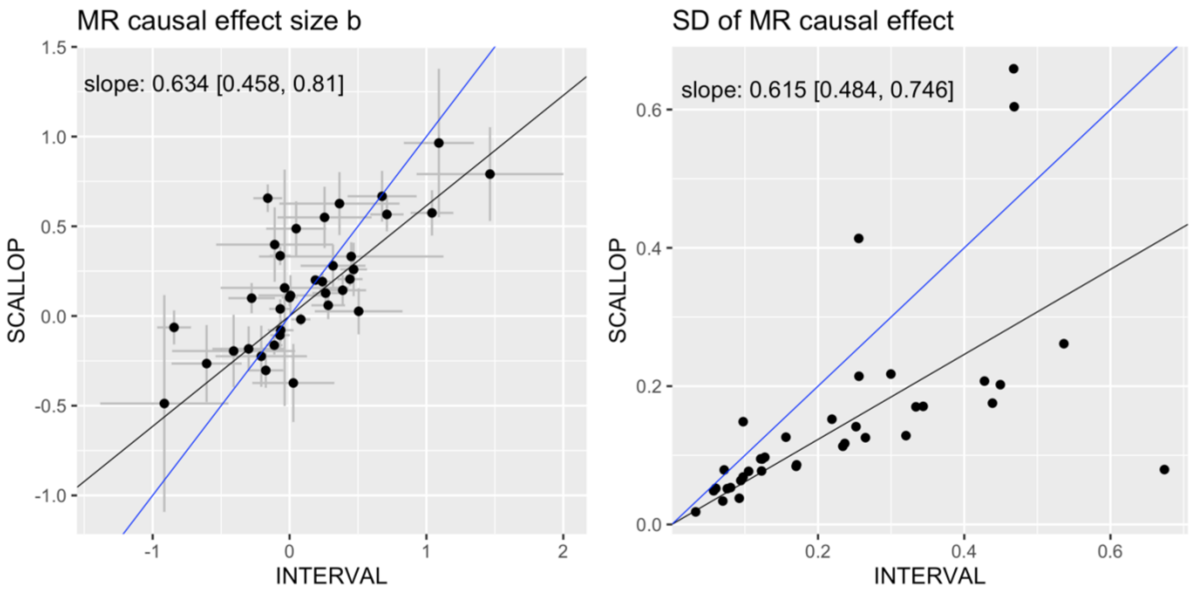

**Supplementary Figure 19.** Transcript-to-protein causal effect comparison between protein (outcome) data from the INTERVAL and SCALLOP consortia. In the left panel, causal MR effect estimates from both datasets are plotted against each other with the error bars being the standard errors of the estimates. In the right panel, the said standard errors are plotted against each other. In both panels, the regression slopes with 95% CI are indicated in the plotting area and plotted are the regression slope (black) and identity (blue) lines.

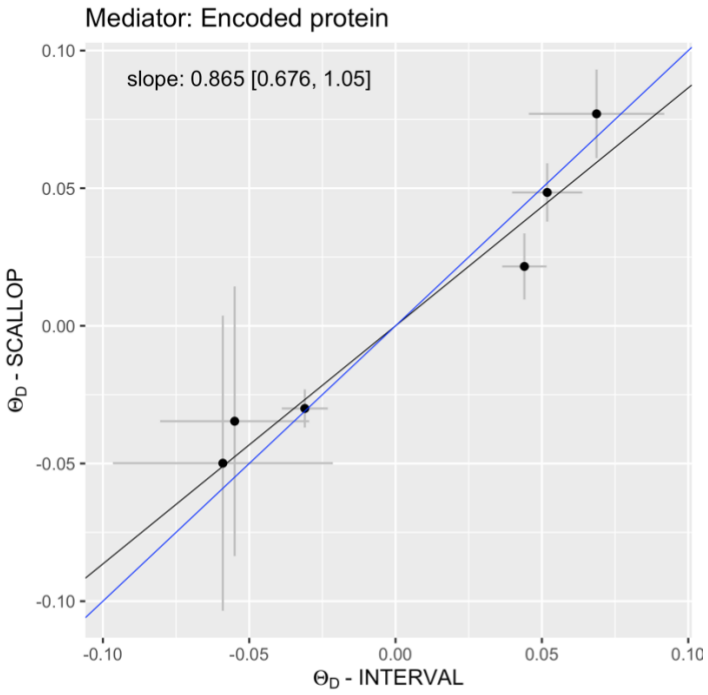

**Supplementary Figure 20.** Validation of the transcript-to-trait mediation analysis through the encoded protein in the SCALLOP consortium. Mediation analyses were conducted through the encoded protein with data once from the INTERVAL and once from the SCALLOP consortia. The resulting direct effects are plotted against each other with the error bars being the standard errors of the estimates. The regression slope with 95% CI is indicated in the plotting area and plotted are the regression slope (black) and identity (blue) lines.

### Multi-omics mechanisms of action

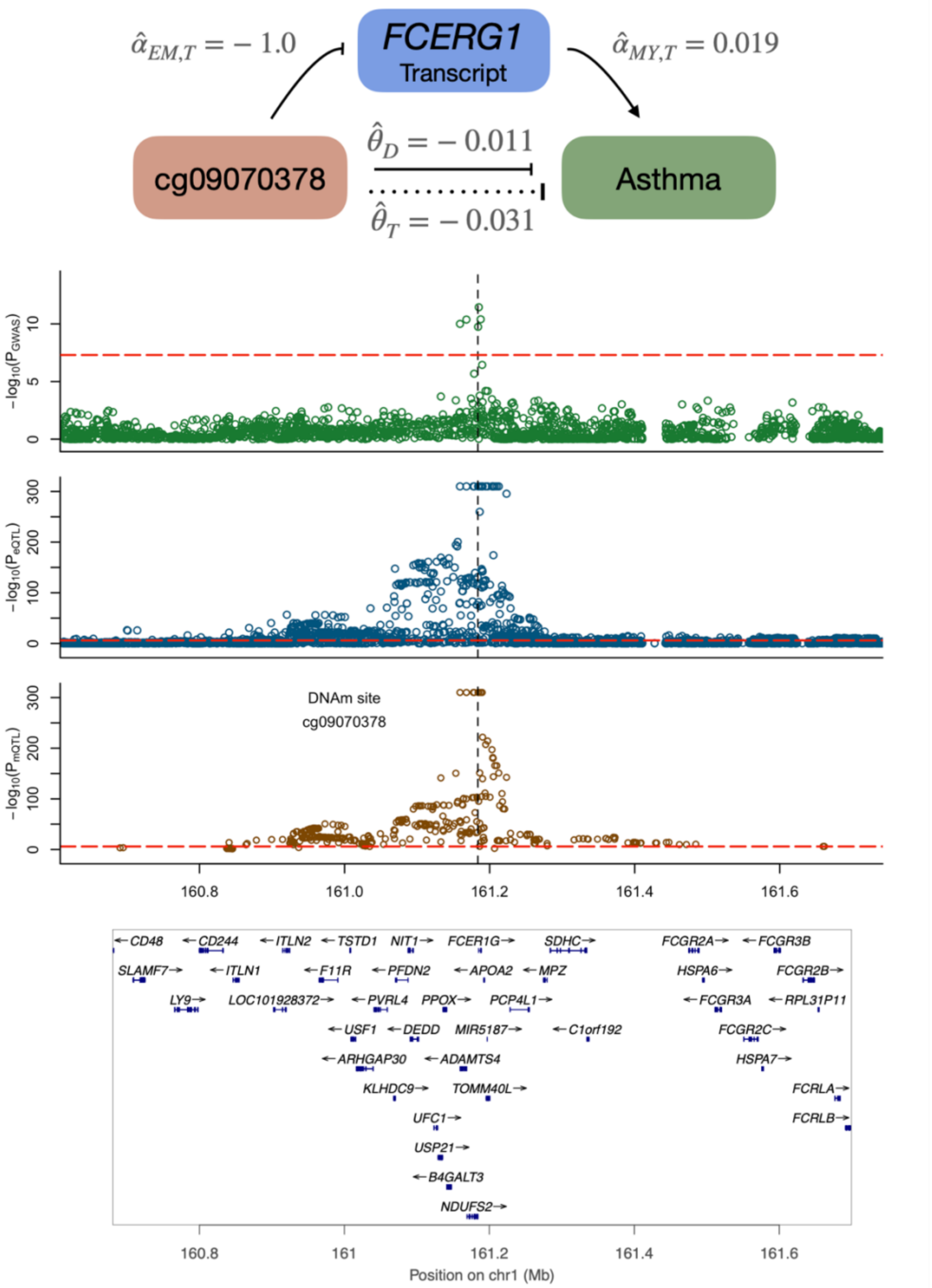

**Supplementary Figure S21**. Plausible DNAm-transcript-trait regulatory mechanism for asthma disease at the *FCERG1* locus. The top row displays a schematic of the mechanism with the calculated univariable and multivariable MR effects. The three following rows show the regional SNP associations (-$\log_{10}$(p-values)) with the trait, transcript and DNAm probe, respectively. Red dashed lines indicate the significance thresholds of the respective SNP associations and the vertical black dashed line represents the DNAm probe position. The bottom row shows the positions of the genes in the locus with their respective strand direction.

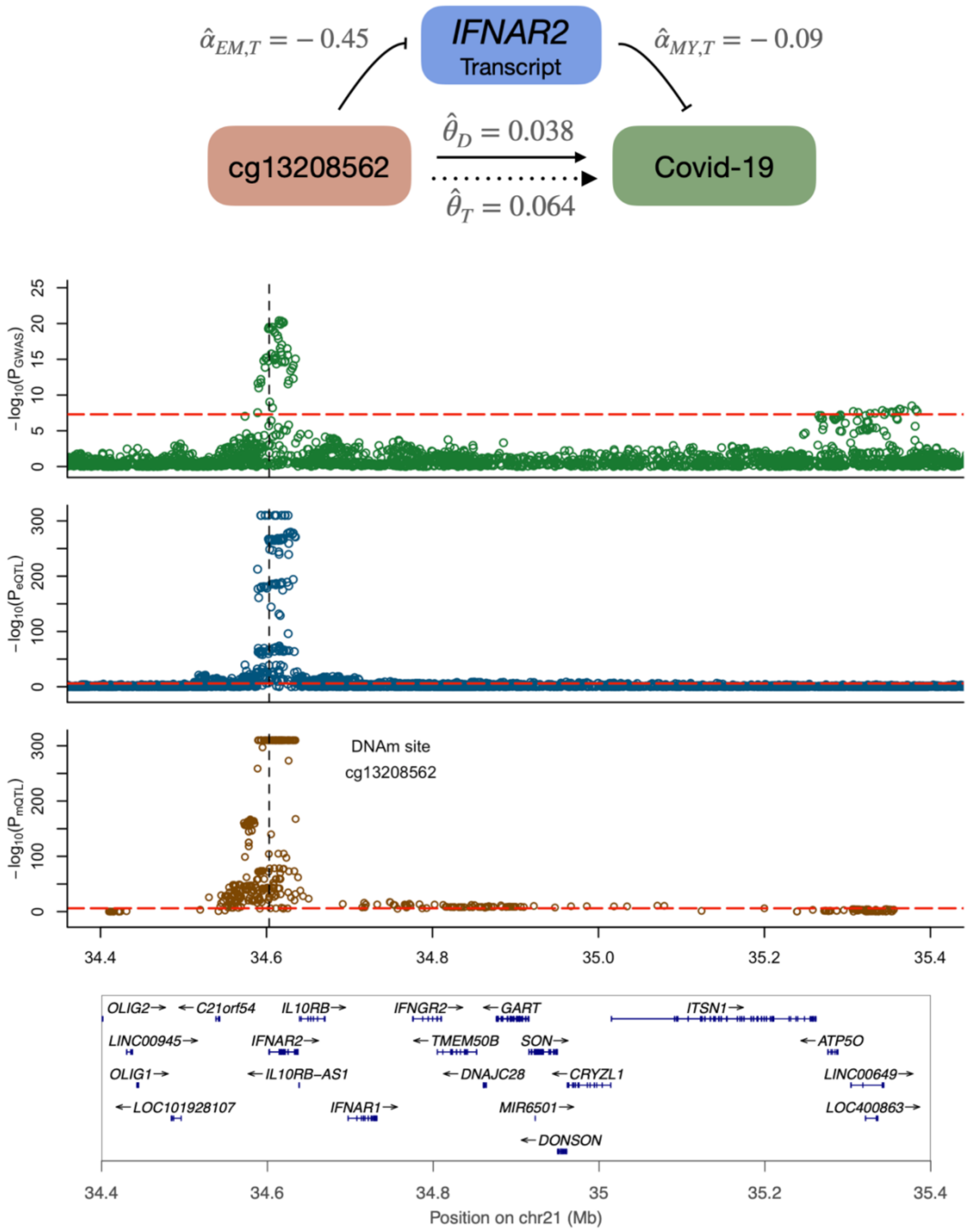

**Supplementary Figure S22.** Plausible DNAm-transcript-trait regulatory mechanism for Covid-19 (hospitalized vs population) at the *IFNAR2* locus. Same figure composition as Figure S21.

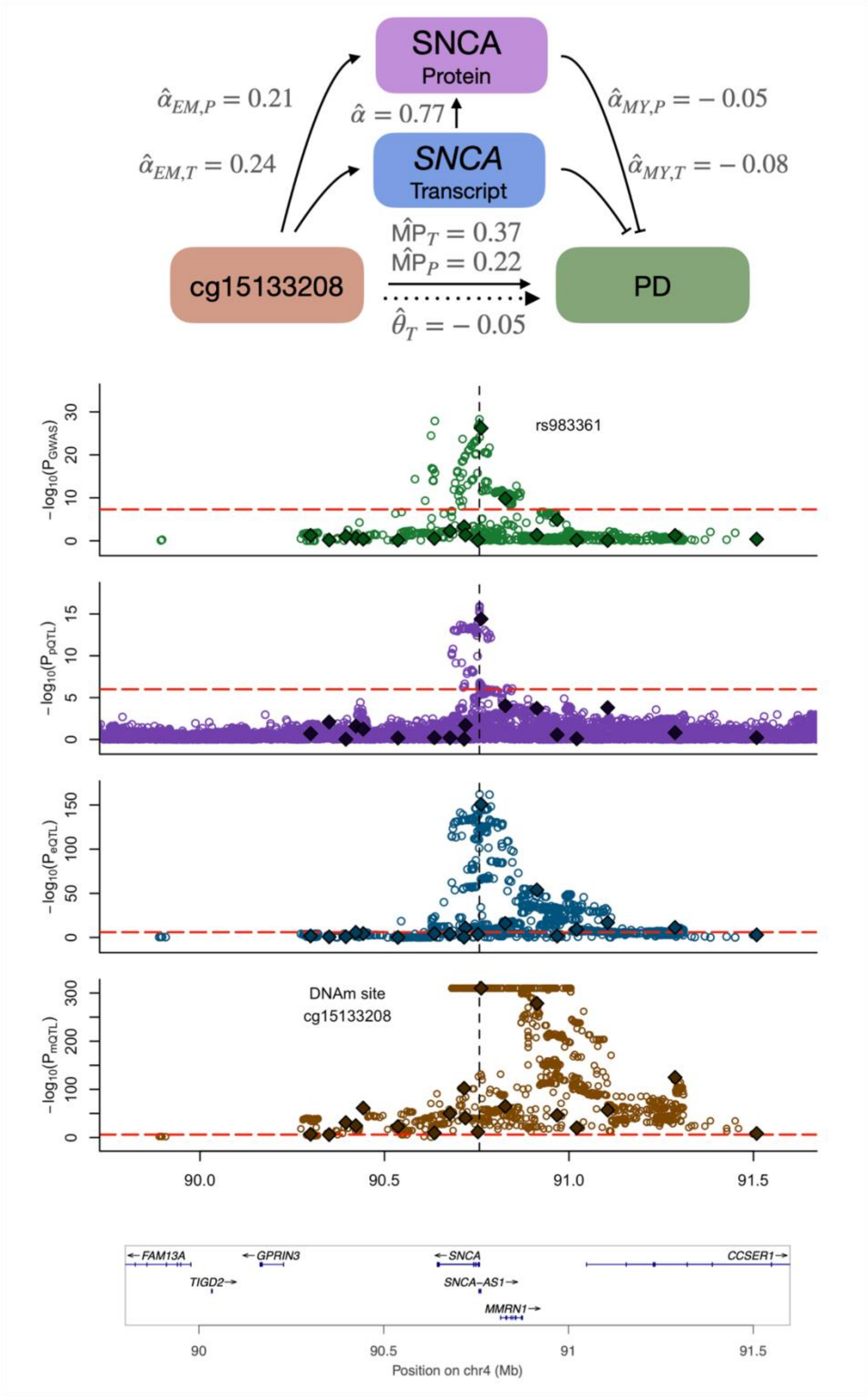

**Supplementary Figure S23**. Plausible DNAm-transcript/protein-trait regulatory mechanisms for Parkinson's disease (PD) at the *SNCA* locus. Same figure composition as Figure 6.

#### Mediation analysis: Technical aspects

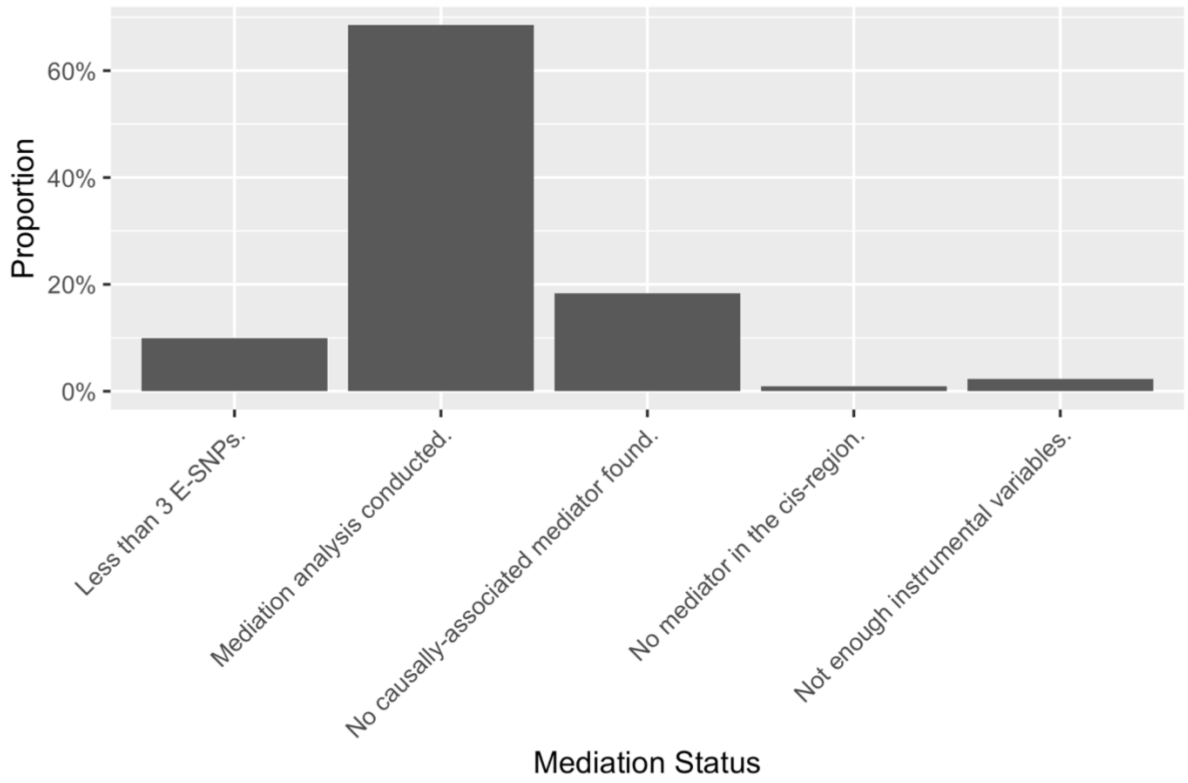

**Supplementary Figure 24.** Distribution of the number of times DNAm-trait pairs could be evaluated in the mediation through transcripts in *cis* ("Mediation analysis conducted") as opposed to no mediation analysis being conducted due to the following reason: "Less than 3 E-SNPs" (= less than 3 exposure-associated SNPs were among all the instrumental variables), "No causally-associated mediator found" (= there were transcripts in the *cis* region, but with no significant DNAm-to-transcript MR effect), "No mediator in the cis-region" (= no transcript was found in a window of 500 kB around the DNAm site) and "Not enough instrumental variables" (= not enough IVs to calculate a direct effect given the number of mediators).

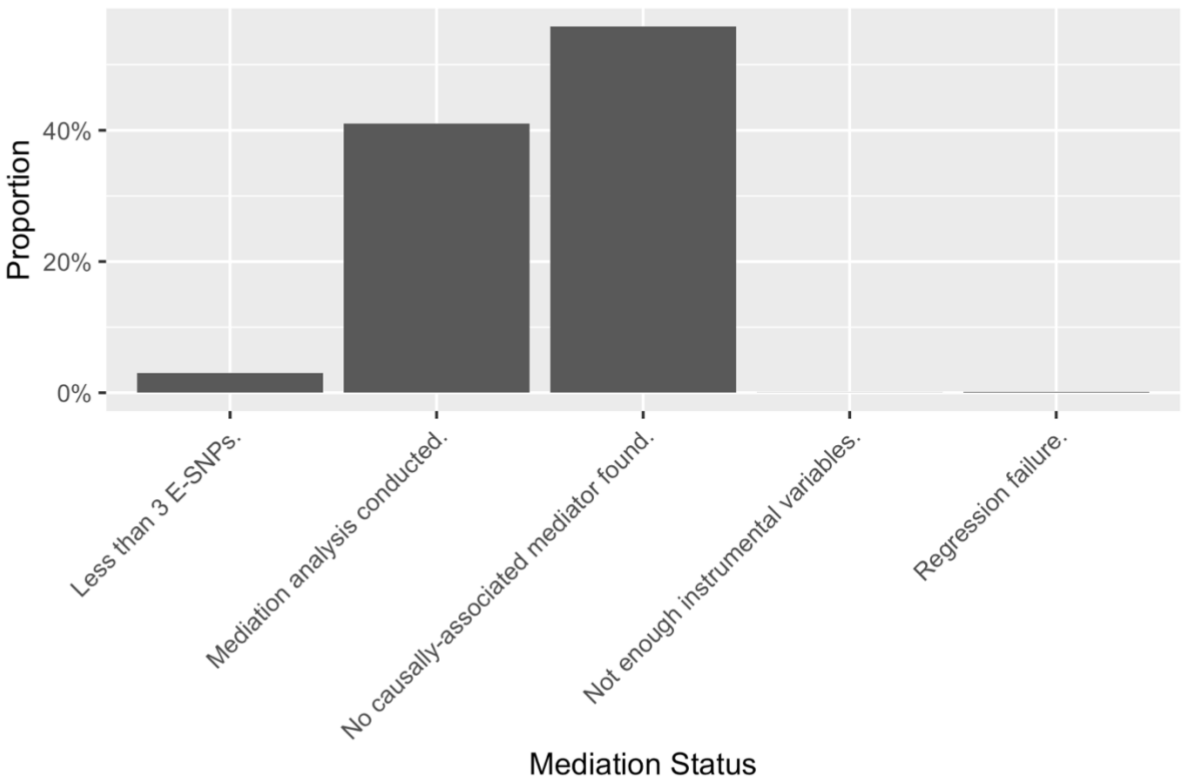

**Supplementary Figure 25.** Distribution of the number of times transcript-trait pairs could be evaluated in the mediation through proteins (encoded ones in addition to causally associated proteins in *trans*; "Mediation analysis conducted") as opposed to no mediation analysis being conducted due to the following reason: "Less than 3 E-SNPs" (= less than 3 exposure-associated SNPs were among all the instrumental variables), "No causally-associated mediator found" (= no encoded protein was present and no additional protein was causally associated to the transcript), "Not enough instrumental variables" (= not enough IVs to calculate a direct effect given the number of mediators) and "Regression failure" (= the regression did not produce numerical estimates likely due to collinearity issues).

### Supplementary Tables

**Supplementary Table 1.** Outcome traits evaluated in the MVMR mediation analyses.

| Abbreviation | Trait | Physiological category | Structural category | Study/ Consortium | Reference | Sample Size N ($N_{cases}$/$N_{controls}$) |
|---|---|---|---|---|---|---|
| Albumin | Albumin | Hepatic | Protein | UK Biobank Phenotype Code: 30600 | http://www.nealelab.is/uk-biobank | 315,268 |
| ALP | Alkaline phosphatase | Musculo-skeletal | Protein | UK Biobank Phenotype Code: 30610 | http://www.nealelab.is/uk-biobank | 344,292 |
| ALT | Alanine amino-transferase | Hepatic | Protein | UK Biobank Phenotype Code: 30620 | http://www.nealelab.is/uk-biobank | 344,136 |
| APO A | Apolipoprotein A | Cardio-vascular | Protein | UK Biobank Phenotype Code: 30630 | http://www.nealelab.is/uk-biobank | 313,387 |
| APO B | Apolipoprotein B | Cardio-vascular | Protein | UK Biobank Phenotype Code: 30640 | http://www.nealelab.is/uk-biobank | 342,590 |
| AST | Aspartate aminotransferase | Hepatic | Protein | UK Biobank Phenotype Code: 30650 | http://www.nealelab.is/uk-biobank | 342,990 |
| Asthma | Asthma | Inflammation | Disease | UK Biobank | Han et al., 2020 [1] | 64,538 / 329,321 |
| Baldness | Baldness | Hormonal | Physical measurement | UK Biobank | Yap et al., 2018 [2] | 205,327 |
| Total bilirubin | Total bilirubin | Hepatic | Metabolite | UK Biobank Phenotype Code: 30840 | http://www.nealelab.is/uk-biobank | 342,829 |
| Bone mineral density | Bone mineral density | Musculo-skeletal | Physical measurement | UK Biobank Phenotype Code: 3148 | http://www.nealelab.is/uk-biobank | 206,496 |
| BMI | Body mass index | Adiposity | Physical measurement | UK Biobank Phenotype Code: 21001 | http://www.nealelab.is/uk-biobank | 359,983 |
| Basal metabolism rate | Basal metabolic rate | Adiposity | Physical measurement | UK Biobank Phenotype Code: 23105 | http://www.nealelab.is/uk-biobank | 354,825 |
| Calcium | Calcium | Cardio-vascular | Metabolite | UK Biobank Phenotype Code: 30680 | http://www.nealelab.is/uk-biobank | 315,153 |
| CAD | Coronary artery diseases | Cardio-vascular | Disease | CARDIoGRAMplusC4D | CARDIoGRAMplusC4D Consortium, 2015 [3] | 60,801 / 123,504 |
| Covid-19, hospitalized | Covid-19, hospitalized vs population | Inflammation | Disease | COVID-19 host genetics initiative, release 6 | https://www.covid19hg.org | 24,274 / 2,061,529 |
| C-reactive protein | C-reactive protein | Inflammation | Protein | UK Biobank Phenotype Code: 30710 | http://www.nealelab.is/uk-biobank | 343,524 |

| | | | | | | |
|---|---|---|---|---|---|---|
| Creatinine | Serum creatinine | Renal | Metabolite | UK Biobank Phenotype Code: 30510 | http://www.nealelab.is/uk-biobank | 350,812 |
| Cystatin C | Cystatin C | Renal | Protein | UK Biobank Phenotype Code: 30720 | http://www.nealelab.is/uk-biobank | 344,264 |
| Diastolic BP | Diastolic blood pressure | Cardio-vascular | Physical measurement | UK Biobank Phenotype Code: 4079 | http://www.nealelab.is/uk-biobank | 340,162 |
| eGFR | Estimated glomerular filtration rate | Renal | Physical measurement | CKDGen | Wuttke et al., 2019 [4] | 567,460 |
| Fluid intelligence | Fluid intelligence | Neuro-psychiatric | Physical measurement | UK Biobank Phenotype Code: 20016 | http://www.nealelab.is/uk-biobank | 117,131 |
| Forced vital capacity | Forced vital capacity | Musculo-skeletal | Physical measurement | UK Biobank Phenotype Code: 20151 | http://www.nealelab.is/uk-biobank | 272,338 |
| GGT | Gamma-glutamyl transferase | Hepatic | Protein | UK Biobank Phenotype Code: 30730 | http://www.nealelab.is/uk-biobank | 344,104 |
| Glucose | Glucose | Diabetes | Metabolite | UK Biobank Phenotype Code: 30740 | http://www.nealelab.is/uk-biobank | 314,916 |
| Grip strength | Hand grip strength | Musculo-skeletal | Physical measurement | UK Biobank Phenotype Code: 47 | http://www.nealelab.is/uk-biobank | 359,729 |
| HbA1c | Glycated hemoglobin | Diabetes | Protein | UK Biobank Phenotype Code: 30750 | http://www.nealelab.is/uk-biobank | 344,182 |
| HDL | High-density lipoprotein cholesterol | Cardio-vascular | Lipid | UK Biobank Phenotype Code: 30760 | http://www.nealelab.is/uk-biobank | 315,133 |
| Height | Height | Musculo-skeletal | Physical measurement | UK Biobank Phenotype Code: 50 | http://www.nealelab.is/uk-biobank | 272,338 |
| IBD | Inflammatory bowel disease | Inflammation | Disease | GWAS meta-analysis | Liu et al., 2015 [5] | 12,882 / 21,770 |
| IGF-1 | Insulin-like growth factor 1 (IGF-1) | Hormonal | Protein | UK Biobank Phenotype Code: 30770 | http://www.nealelab.is/uk-biobank | 342,439 |
| Alzheimer's disease | Late onset Alzheimer's disease | Neuro-psychiatric | Disease | GWAS meta-analysis | Lambert et al., 2013 [6] | 17,008 / 37,154 |
| LDL | Low-density lipoprotein cholesterol | Cardio-vascular | Lipid | UK Biobank Phenotype Code: 30780 | http://www.nealelab.is/uk-biobank | 343,621 |
| Lymphocyte count | Lymphocyte count | Hemato-logical | Lipid | UK Biobank Phenotype Code: 30120 | http://www.nealelab.is/uk-biobank | 349,856 |
| Parkinson's disease | Parkinson's disease | Neuro-psychiatric | Disease | GWAS meta-analysis | Nalls et al., 2019 [7] | 56,300 / 1,400,000 |
| Platelet count | Platelet count | Hemato-logical | Physical measurement | UK Biobank Phenotype Code: 30080 | http://www.nealelab.is/uk-biobank | 350,474 |

| | | | | | | |
|---|---|---|---|---|---|---|
| Pulse rate | Pulse rate | Cardio-vascular | Physical measurement | UK Biobank Phenotype Code: 102 | http://www.nealelab.is/uk-biobank | 340,162 |
| Rheumatoid arthritis | Rheumatoid arrthritis | Inflammation | Disease | GWAS meta-analysis | Okada et al., 2013 [8] | 19,234 / 61,565 |
| Red blood cell count | Red blood cell count | Hemato-logical | Physical measurement | UK Biobank Phenotype Code: 30010 | http://www.nealelab.is/uk-biobank | 350,475 |
| Systolic BP | Systolic blood pressure | Cardio-vascular | Physical measurement | UK Biobank Phenotype Code: 4080 | http://www.nealelab.is/uk-biobank | 340,159 |
| Schizophrenia | Schizophrenia | Neuro-psychiatric | Disease | Psychiatric Genomics Consortium | Schizophrenia Working Group of the Psychiatric Genomics Consortium [9] | 36,989 / 113,075 |
| SHBG | Sex hormone-binding globulin | Hormonal | Protein | UK Biobank Phenotype Code: 30830 | http://www.nealelab.is/uk-biobank | 312,215 |
| Stroke | Stroke | Cardio-vascular | Disease | MEGASTROKE Consortium | Malik et al., 2018 [10] | 40,585 / 406,111 |
| Total cholesterol | Total cholesterol | Cardio-vascular | Lipid | UK Biobank Phenotype Code: 30690 | http://www.nealelab.is/uk-biobank | 344,278 |
| T2D | Type 2 diabetes | Diabetes | Disease | DIAGRAM Consortium & UK Biobank | Mahajan et al., 2018 [11] | 74,124 / 824,006 |
| Testosterone | Testosterone | Hormonal | Lipid | UK Biobank Phenotype Code: 30850 | http://www.nealelab.is/uk-biobank | 312,102 |
| Triglycerides | Triglycerides | Cardio-vascular | Lipid | UK Biobank Phenotype Code: 30870 | http://www.nealelab.is/uk-biobank | 343,992 |
| Urate | Serum urate | Renal | Metabolite | UK Biobank Phenotype Code: 30880 | http://www.nealelab.is/uk-biobank | 343,836 |
| Urea | Serum urea | Renal | Metabolite | UK Biobank Phenotype Code: 30670 | http://www.nealelab.is/uk-biobank | 344,052 |
| Vitamin D | Vitamin D | Musculo-skeletal | Metabolite | UK Biobank Phenotype Code: 30890 | http://www.nealelab.is/uk-biobank | 329,247 |
| WHR adj. BMI | Waist-to-hip ratio adjusted for BMI | Adiposity | Physical measurement | GIANT & UK Biobank | Pulit et al., 2019 [12] | 694,649 |

**Supplementary Table 2.** Default simulation parameters to mimic the setting of DNAm or transcript levels as exposure.

| Parameter | DNAm-exposure | Transcript level-exposure |
|---|---|---|
| $N_E$ | 30,000 | 30,000 |
| $N_M$ | 30,000 | 3,000 |
| $N_Y$ | 300,000 | 300,000 |
| $N_{med}$ | 20 | 500 |
| $N_{med,sig}$ | 3 | 2 |
| $m_E$ | 6 | 20 |
| $m_M$ | 3 | 3 |
| $h^2_E$ | 0.35 | 0.2 |
| $h^2_{M,low}$ | 0.005 | 0.015 |
| $h^2_{M,lhigh}$ | 0.02 | 0.08 |
| $var(\alpha^{EM})$ | 0.1 | 0.15 |
| $var(\alpha^{MY})$ | 0.0064 | 0.0007 |
| ρ | 0.15 | 0.15 |
| MP | 0.3 | 0.15 |
| $P_{EM}$ | 0.01 | 0.001 |

**Supplementary Table 3.** Number of exposure-trait pairs that could be evaluated in the three mediation analyses 1) DNAm→trait through transcripts, 2) DNAm→trait through proteins, 3) transcript→trait through proteins and in each of the following steps: "All" (=all pairs with a significant exposure-to-trait causal effect), "With testable mediators" (=pairs with at least 1 testable mediator in the *cis*-region if DNAm was the exposure; always the same as "All" in the transcript-exposure setting since proteins in *trans* were assessed as mediators), "With causally-associated mediator" (=pairs with at least 1 causally-associated mediator), "Successful mediation" (=pairs for which the mediation analysis through at least 1 mediator succeeded with potential failures being shown in Figure S24 and S25), and "Successful mediaton through focal mediator" (=pairs for which a mediation through a focal mediator could be conducted which was the "top" transcript/protein in the DNAm-exposure setting and the encoded protein in the transcript-exposure setting). Note that the number of pairs in the "All" step slightly vary between the DNAm-to-trait via transcripts and proteins analyses due to the data harmonization step. In theory, both numbers should be the same, but given that some SNPs were not present in all datasets, there were insufficient exposure IVs for a few DNAm probes.

| Mediation setting | All | With testable mediators | With causally-associated mediator | Successful mediation | Successful mediation through focal mediator |
|---|---|---|---|---|---|
| **DNAm-to-trait via transcripts** | 3,020 | 2,992 | 2,438 | 2,069 | 2,069 |
| **DNAm-to-trait via proteins** | 3,048 | 2,388 | 336 | 328 | 328 |
| **Transcript-to-trait via proteins** | 3,848 | 3,848 | 1,701 | 1,577 | 333 |

**Supplementary Table 4.** Enrichment analysis of negative DNAm-to-transcript causal effects within each annotation group (i.e. DNAm probe annotations with respect to the assessed transcript). The first two columns show the number of distinct DNAm-transcript pairs with negative and positive causal effects, respectively. An enrichment analysis for negative causal effects was conducted as a two-sided Fisher's test where each annotation group was tested against the remaining other groups combined. Annotation groups significantly enriched or deprived for negative causal effects (after correcting for multiple testing at P < 0.05/6) are highlighted in bold.

| Annotation group | DNAm → Transcript negative effect | DNAm → Transcript positive effect | Proportion of negative effects | OR (negative effect enrichment) | P-value (negative effect enrichment) |
|---|---|---|---|---|---|
| **1stExon** | 291 | 188 | 0.608 | **1.33** | **2.67E-03** |
| 3'UTR | 863 | 828 | 0.510 | 0.89 | 1.63E-02 |
| 5'UTR | 1773 | 1429 | 0.554 | 1.07 | 8.03E-02 |
| **Body** | 9675 | 8824 | 0.523 | **0.87** | **2.15E-10** |
| **TSS1500** | 4380 | 3433 | 0.561 | **1.12** | **1.24E-05** |
| **TSS200** | 1702 | 1284 | 0.570 | **1.15** | **3.81E-04** |
